# Deep learning-driven imaging of cell division and cell growth across an entire eukaryotic life cycle

**DOI:** 10.1101/2024.04.25.591211

**Authors:** Shreya Ramakanth, Taylor Kennedy, Berk Yalcinkaya, Sandhya Neupane, Nika Tadic, Nicolas E. Buchler, Orlando Argüello-Miranda

## Abstract

The life cycle of biomedical and agriculturally relevant eukaryotic microorganisms involves complex transitions between proliferative and non-proliferative states such as dormancy, mating, meiosis, and cell division. New drugs, pesticides, and vaccines can be created by targeting specific life cycle stages of parasites and pathogens. However, defining the structure of a microbial life cycle often relies on partial observations that are theoretically assembled in an ideal life cycle path. To create a more quantitative approach to studying complete eukaryotic life cycles, we generated a deep learning-driven imaging framework to track microorganisms across sexually reproducing generations. Our approach combines microfluidic culturing, life cycle stage-specific segmentation of microscopy images using convolutional neural networks, and a novel cell tracking algorithm, FIEST, based on enhancing the overlap of single cell masks in consecutive images through deep learning video frame interpolation. As proof of principle, we used this approach to quantitatively image and compare cell growth and cell cycle regulation across the sexual life cycle of *Saccharomyces cerevisiae*. We developed a fluorescent reporter system based on a fluorescently labeled Whi5 protein, the yeast analog of mammalian Rb, and a new High-Cdk1 activity sensor, LiCHI, designed to report during DNA replication, mitosis, meiotic homologous recombination, meiosis I, and meiosis II. We found that cell growth preceded the exit from non-proliferative states such as mitotic G1, pre-meiotic G1, and the G0 spore state during germination. A decrease in the total cell concentration of Whi5 characterized the exit from non-proliferative states, which is consistent with a Whi5 dilution model. The nuclear accumulation of Whi5 was developmentally regulated, being at its highest during meiotic exit and spore formation. The temporal coordination of cell division and growth was not significantly different across three sexually reproducing generations. Our framework could be used to quantitatively characterize other single-cell eukaryotic life cycles that remain incompletely described. An off-the-shelf user interface *Yeastvision* provides free access to our image processing and single-cell tracking algorithms.

## Introduction

A quantitative study of the microbial life cycle is crucial to understand ecological and evolutionary processes that can be used to control parasites and pathogens ^1–4^. However, defining the structure of a microbial life cycle traditionally relies on partial microscopy observations or genetic and theoretical evidence ^5^. Although biochemical methods, such as immunostainings or single cell DNA/RNA sequencing, can provide molecular descriptions of developmental transitions ^3, 6–9^ , only live-cell imaging can directly visualize entire microbial life cycles in single cells *in vivo,* as demonstrated for bacterial ^10, 11^ and eukaryotic ^12^ life cycles. However, quantitative life cycle imaging remains a challenge for sexually reproducing microbes, many of which are important human parasites ^3, 13–15^. This difficulty arises from the need to track single cells through various morphological transitions of sexual reproduction, including the creation of haploid sex cells through meiosis, the reconstitution of diploidy through mating, and other changes in cell size and ploidy ^16^. The lack of standardized methods to continuously visualize complete life cycles has led, in extreme cases, to confuse morphologically diverse life cycle stages as different species ^17, 18^. Furthermore, even though genetic evidence like DNA recombination rates indicate the presence of meiosis, the existence of sexual life cycles in some microorganisms remains uncertain ^19, 20^. In this work, we provide a solution to quantitatively study entire microbial sexual life cycles by using a combination of microfluidic manipulations, long-term live cell imaging, and deep learning algorithms for cell detection-tracking across a series of time-lapse images.

As proof of principle, we image the sexual life cycle of a model microorganism, the yeast *Saccharomyces cerevisiae*. In the wild, *S. cerevisiae* cells are diploids that proliferate in nutrient-rich conditions using mitotic cell division, whereby one round of DNA replication follows one round of chromosome segregation and cell division ^21^ (**Fig 1 A**, diploid cell cycle). Transitions through cell growth and cell division are driven by the activation of the cycling dependent kinase Cdk1; periods of low Cdk1 activity are non-proliferative states, whereas periods of high Cdk1 activity promote DNA replication and mitosis or meiosis ^21, 22^. Cells challenged by starvation will arrest their proliferation and enter protective dormant states through quiescence or meiosis (**Fig 1 A**, starvation stress response, arrest, quiescence, meiosis). During quiescence, cells become stress resistant due to the reorganization of cell wall, metabolic, organelle, and gene expression patterns, which are reversible allowing cells to re-engage cell cycle under rich conditions ^23–26^. By contrast, during meiosis, the genome is irreversibly haploidized through a specialized round of DNA replication that is followed by homologous chromosome recombination and two rounds of chromosome segregation that lead to the formation of four haploid nuclei (**Fig 1 A**, meiosis I, meiosis II) ^16^. Healthy meiotic divisions lead to the production of four haploid nuclei that are individually packaged into highly stress-resistant spores surrounded by the membrane and cell wall of the former cell, or ascus ^27, 28^ (**Fig 1 A**, sporulation, ascus). Germinating spores can be of two opposite mating types, MAT a or MAT alpha, which can directly mate inside the ascus during germination to restore diploidy (**Fig 1 A**, ascus mating) ^29, 30^. Wild-type (WT) yeast spores are homothallic, or able to change their mating type to ensure mating with adjacent cells, even between mother-daughter or sister-sister cells (**Fig 1 A**, Homothallic, inbreeding). Laboratory strains, on the other hand, are often heterothallic; cells cannot switch their mating types, if mating inside the ascus fails, cells will undergo mating only until finding opposite mating type cells (**Fig 1 A,** heterothallic, outbreeding).^29, 30^

**Fig 1.**
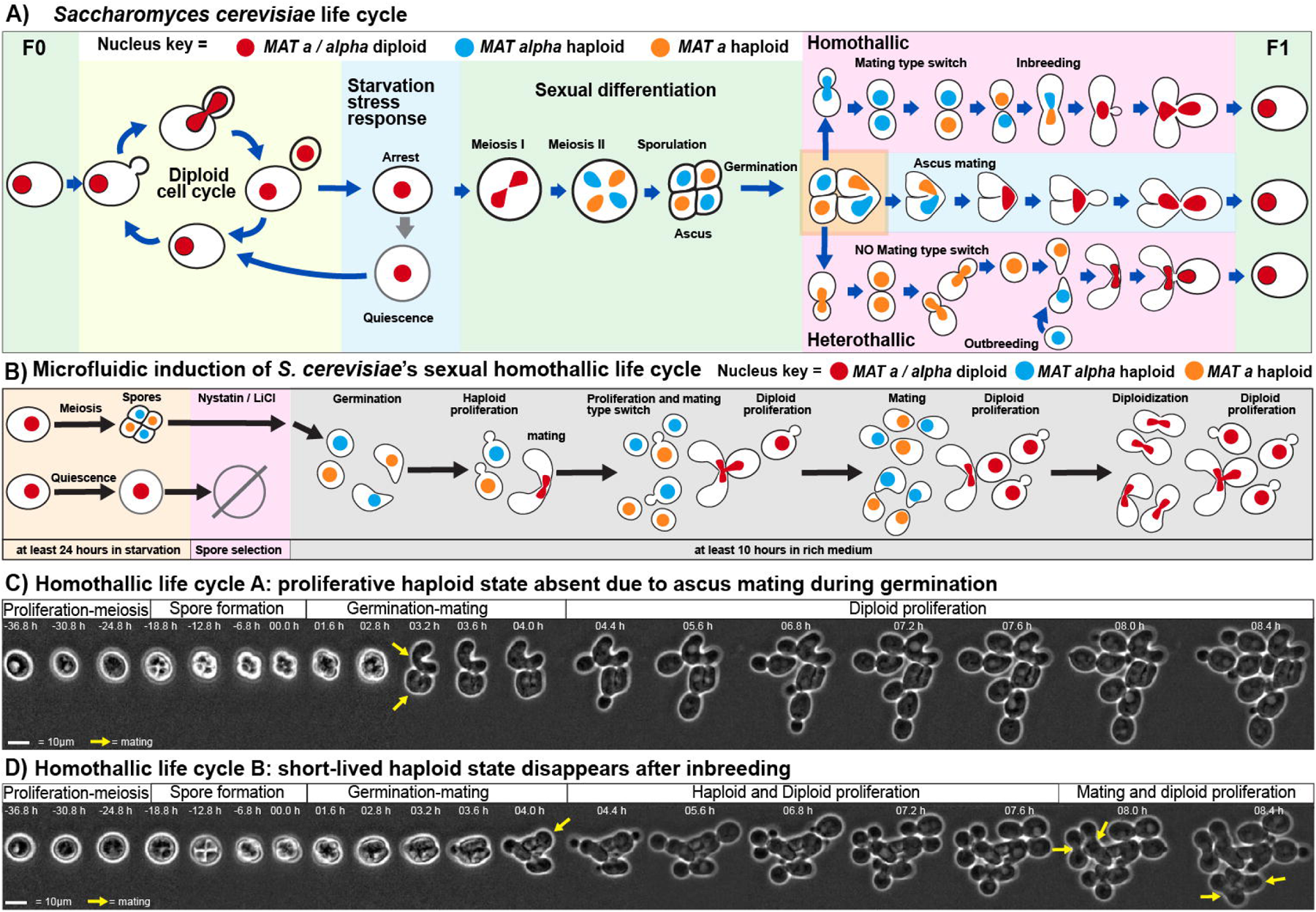
Microfluidics life cycle induction in the model eukaryote *Saccharomyces cerevisiae*. **(A)** Schematic of *S. cerevisiae* life cycle. F0 = parental generation, F1 = filial generation 1 **(B)** Schematic microfluidic protocol to specifically trigger *S. cerevisiae’s* sexual life cycle by selecting against non-sporulated cells that can outcompete germinating spores. Nuclei represented as solid shapes inside the cells. **(C-D)** Representative time lapse micrographs of computationally aligned yeast cells undergoing **(C)** homothallic life cycle type A, in which sporulation is followed by ascus mating directly leading to two diploid cells without an intervening proliferative haploid state, or **(D)** Homothallic life cycle type B, in which sporulation is followed by partial ascus mating producing one diploid and two proliferative haploid cells which eventually diploidize through inbreeding.

The activity of the Cdk1 kinase regulates many life cycle transitions via phosphorylation of cell cycle and meiosis regulators ^31, 32^. Low-Cdk1 activity states include mitotic G1, proliferation arrest in response to mating pheromone ^33^ or nutritional or physical stress ^34^, pre-meiotic G1 ^35^, quiescence ^36^, and the spore state^28^,. States with increased Cdk1 kinase activity allow for DNA replication and cell division, referred to as High-Cdk1 states, which can be mitotic or meiotic ^37, 38^. The transitions between Low-and High-Cdk1 states are regulated by protein regulatory networks known as checkpoints ^39–41^. For instance, newborn cells remain in the low Cdk1-Activity state of G1 until they cross a size threshold required to enter the higher Cdk1 kinase state for DNA replication and mitosis^42^. In yeast, this G1-to-S-phase transition is known as the START checkpoint ^43^. The Low Cdk1 kinase state before START is maintained by G1 stabilizers, which include APC/C-Cdh1 ubiquitination-mediated proteolysis of M-phase proteins and Cdk1 activators ^44, 45^, the accumulation of Cdk1 inhibitors such as Sic1 ^46, 47^, and the transcriptional repression of the mitotic program by Whi5 ^48^, the yeast functional homolog of the mammalian Rb tumor suppressor. To trigger G1 exit, cell growth dilutes Whi5 allowing the expression of G1/S cyclins ^49^ ^42^, which phosphorylate and further inactivate Whi5, triggering positive feedback loops that inactivate the remaining key G1 stabilizers ^50^.

*S. cerevisiae’s* life cycle transitions between low-and high-Cdk1 states can be quantitatively imaged by integrating microfluidic culturing with timelapse microscopy of strains expressing fluorescent cell cycle reporters ^51, 52^, an approach also used in other microorganisms. However, the biggest bottleneck for quantitatively imaging full life cycles is the lack of robust computational methods to detect and track single cells through morphologically complex or spatially dynamic transitions, such as mitotic exponential proliferation, spore release during germination, and cell fusion while mating ^53, 54^. Although recent deep-learning image processing approaches based on convolutional neural networks can segment and track single yeast cells in microscopy images, most methods are predominantly aimed at mitotic proliferation, which does not display the morphological variation of life cycle stages such as sporulation, germination, and mating ^55–59^.

In this work, we provide a solution to track single microbial cells through their entire sexual life cycle using Frame Interpolation Enhanced Single-cell Tracking (FIEST), a computational framework that leverages a life cycle cell segmentation approach based on Cellpose models ^60, 61^, in combination with frame interpolation using real-time flow estimation (RIFE) ^62^. We use this cell tracking method to quantitatively visualize Cdk1 activity states throughout *S. cerevisiae*’s full life cycle via a fluorescently-tagged Whi5 reporter and a fluorescent <u>Li</u>fe <u>C</u>ycle <u>H</u>igh-Cdk1 activity Indicator (LiCHI) for both mitotic and meiotic High-Cdk1 states. Using this approach, we showed that Whi5 nuclear accumulation differs greatly across developmental stages within a single life cycle, and this difference persists among ancestor and descendant cells across sexually reproducing generations. We show that the exit from non-proliferative states such as mitotic G1, pre-meiotic G1, and the G0 spore state, is characterized by a decreased in the total cell concentration of Whi5, which is consistent with a Whi5 dilution model to trigger exit from non-proliferative states ^63^. The FIEST pipeline is available as an off-the-shelf graphics user interface (GUI) that provides access to our pretrained models, single cell tracking algorithms, and video frame interpolation models for the analysis of images and videos of microbial life cycles.

### Microfluidics life cycle induction in the model eukaryote *Saccharomyces cerevisiae*

We first designed a microfluidics protocol to trigger the sexual yeast life cycle based on mimicking environmental nutritional challenges ^27, 64–66^. Briefly, unsynchronized proliferating Wild-type (WT) SK1 diploid cells (F0, or parental generation in Mendelian nomenclature) were loaded in a high-aspect ratio microfluidic device ^67^, and exposed to starvation medium for at least 24 hours to stop proliferation and trigger quiescence or sporulation (**Fig 1 B**). Quiescent and non-sporulated cells were killed by exposure to a 4 M LiCl solution containing 0.3 mg/mL nystatin for one hour followed by distilled water for one hour. This step enriched for sporulated cells and suppressed competition from quiescent cells upon return to rich medium (**Fig S1 A, Video S1**).

The germinating spores displayed two different types of homothallic lifecycle. In the homothallic type A life cycle, mating immediately occurred in the ascus during germination producing the next diploid generation without an intervening proliferative haploid state (**Fig 1 C**), and preventing mating between spores from different asci **(Fig S1 B).** In the homothallic type B life cycle, germinating spores gave rise to proliferative haploid cells that switched mating types to undergo inbreeding (**Fig 1 D**). Other life cycle variations were detected at 12 ± 3 % (n=9), including the production of three (tryads, **Fig S2 C**) or two (dyads, **Fig S2 D**)-spored asci, and infrequent pseudohyphal cells. Our protocol also recapitulated the heterothallic life cycle in laboratory W303 strains that are unable to switch mating type and can proliferate as haploids after germination with minimal mating **(Fig S2 E)**.

We concluded that our microfluidics protocol recapitulated the sexual life cycle of yeast. We focused our experiments on the homothallic life cycle, which had the advantage of rapidly generating the next diploid generation.

### Life cycle stage-specific segmentation

Imaging the full yeast life cycle revealed significant morphological and optical diversity during sporulation (**Fig 2 A**), germination (**Fig 2 B**) and mating (**Fig 2 C**). Current yeast detection algorithms, such as YeastNET ^58^, YeaZ ^55^, BABY ^68^, Cellpose ^60^, and the yeast compatible cell AC/DC functionalities ^69^, however, do not specifically segment cells according to life cycle stage, and often depend on pixel intensity thresholds that can change due to morphological and optical heterogeneity. To overcome this challenge, we leveraged solutions for pixel intensity- and morphology-independent image segmentation developed for human silhouette recognition ^70^ and bacterial and mammalian cell biology ^71, 72^ based on the openpose and cellpose architectures.

**Fig 2.**
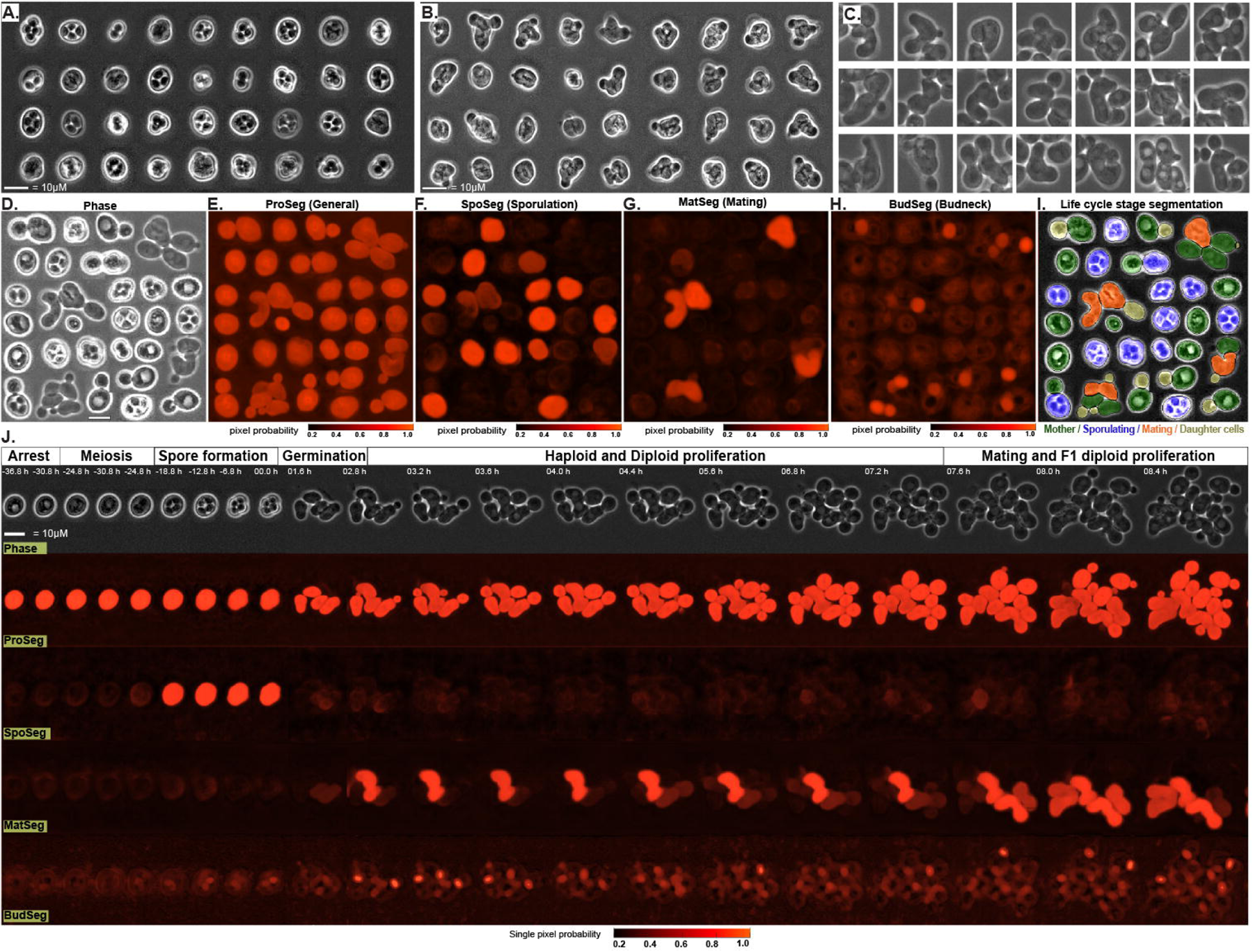
Life cycle stage-specific segmentation enables the detection of morphologically diverse stages in *S. cerevisiae*’s sexual life cycle. **(A-C)** Computationally aligned yeast cells displaying the morphological and optical diversity during the stages of **(A)** sporulation **(B)** germination and **(C)** mating (square side = 25 µm). **(D)** Computationally arranged collage of assorted life cycle stages. Scale bar = 10 µm. **(E-H)** Segmentation of the collage in D as single pixel probability heatmaps using: **(E)** the general segmentation model for detection of yeast cells, ProSeg, **(F)** the sporulation-specific segmentation model, SpoSeg, **(G)** the mating-specific segmentation model, MatSeg, **(H)** the budneck-specific segmentation model, BudSeg, which labels the budneck connection between mother and daughter cells. **(I)** Detection of mother, daughter, sporulating, and mating cells from the collage in D using the results from life cycle-stage specific segmentation models. **(J)** Example segmentation of time series of micrographs with computationally aligned yeast cells undergoing a full sexual life cycle; bottom, time series displaying ProSeg, SpoSeg, MatSeg, and BudSeg segmentation results as single pixel cell probability heatmaps.

To generate life cycle stage-specific segmentations of yeast cells, we trained a series of cellpose models on hundreds of phase contrast images of yeast cells at different life cycle transitions (**Fig 2 D**) and under biochemical or genetic perturbations aimed to increase the phenotypic variability in the training data set such as I.) Proliferation after cell cycle synchronization or under checkpoint-activating drugs II.) Cell size perturbations by deletion of G1 cyclins (*CLN3*) ^73^ or ribosome biogenesis regulators (*SFP1*) ^74^; III.) Cell shape perturbation by overexpression of the APC/C activator Hct1/Cdh1 ^75^; IV.) Organelle perturbations by deletion of vacuole/lysosome inheritance factor (*VAC17*) ^76^; V.) Cell culture conditions at multiple densities and different nutrient depletions. To directly visualize the segmentation models performance, in this work we show the single pixel cell probability heatmap output from the models or the resulting masks derived following the cellpose mask generation methods **(Fig S2 A)**.

The training process resulted in four label-free segmentation models: ProSeg, for general detection of yeast cells across all life cycle stages (**Fig 2 E**), SpoSeg for segmenting cells undergoing sporulation or early germination (**Fig 2 F**), MatSeg for segmenting mating cells (**Fig 2 G**), and BudSeg for segmenting the connection between mother and daughter cells, known as budneck (**Fig 2 H**). Life cycle specific models were obtained through iterative training on human labeled data sets, whereas Budseg was obtained using cells bearing a fluorescent septin ring component, Cdc10 ^77^, to automatically label budneck pixels in phase images **(Fig S2 B,C)**.

To segment yeast cells micrographs containing multiple life cycle stages, we sequentially used ProSeg, MatSeg, SpoSeg, and BudSeg to obtain four detections from the same image creating cell masks for specific life cycle stages (**Fig 2 I, Video S2**). Our models generated cell masks with > 90 % average precision (AP) when compared against human-labeled data (**Table S4),** outperformed pre-existing CNN-segmentation approaches when detecting complex yeast morphology or dense cultures **(Fig S2 D-E, Table S5)**, and enabled the detection of life cycle-specific stages in long term image time series (**Fig 3 J**).

**Fig 3.**
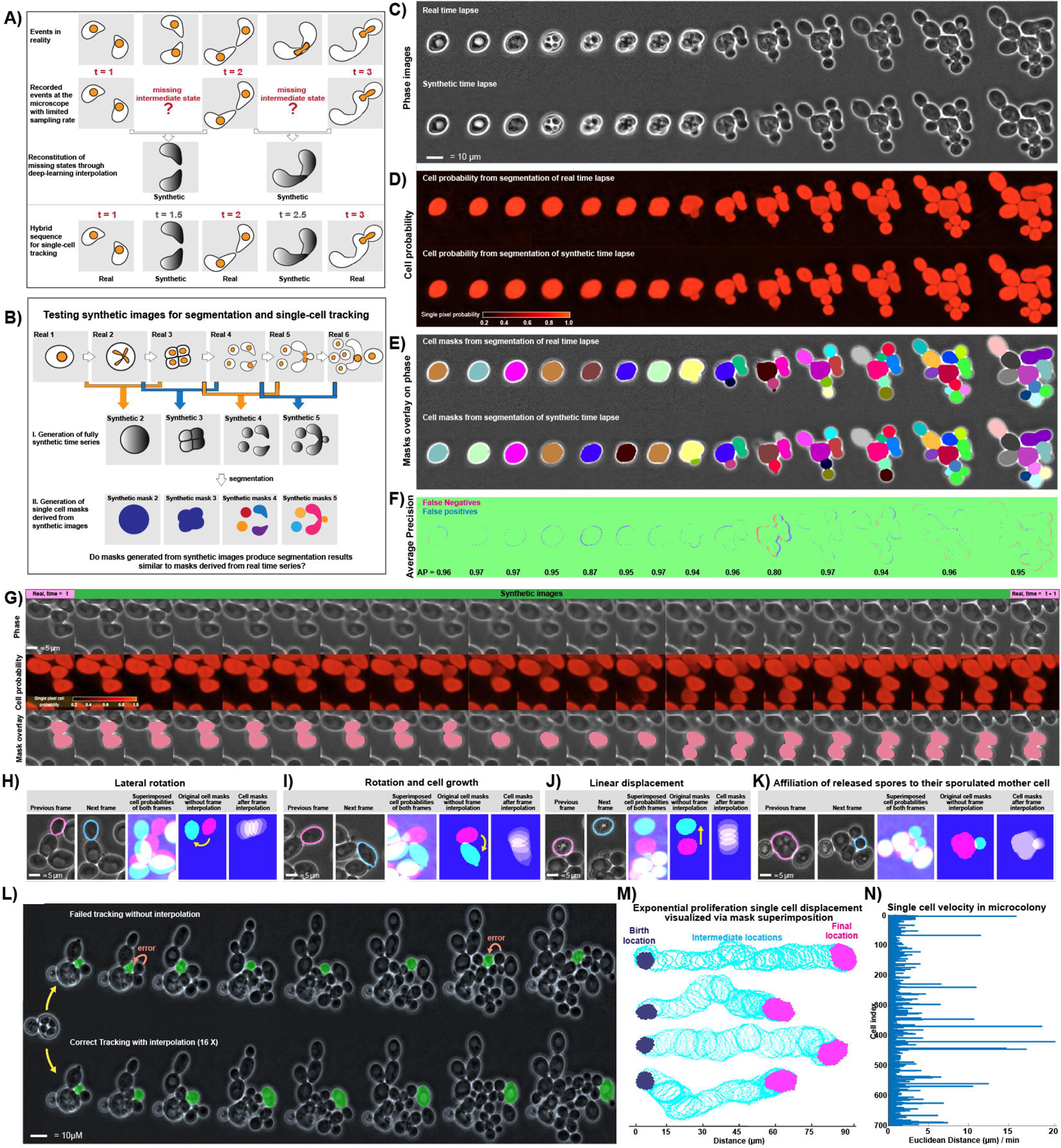
Single cell tracking across *S. cerevisiae*’s life cycle using Frame Interpolation Enhanced Single-cell Tracking (FIEST). **(A)** Schematic of deep learning frame interpolation as a tool to produce potential intermediate cellular states not captured during time lapse microscopy. **(B)** Schematics of the segmentation test for synthetic images. Odd and even frames are interpolated to produce a fully synthetic time series for segmentation comparison against real images. (**C**) Real (top) and synthetic (bottom) time series of concatenated micrographs with computationally aligned yeast cells undergoing a full life cycle. **(D)** Segmentation results as probability heatmaps for the real (top) and synthetic (bottom) time series depicted in C. **(E)** Single cell masks obtained from the real (top) or synthetic (bottom) time series depicted in C as overlay on the phase image. **(F)** Accuracy of cell masks derived from synthetic images compared against masks derived from real images, measured as average precision (AP). Falsely detected pixels, blue, wrongly ignored pixels, red. Bottom values: AP for masks in frame. **(G)** Representative interpolated time series displaying the segmentation and tracking results for a hybrid time series composed of only two real (first and last frames) and synthetic interpolated images of a mating event with abrupt cell displacement. Top, phase images; middle, segmentation results as single pixel probability heatmaps; Bottom, tracked mask for the mating cells across real and interpolated images. **(H-J)** Schematic examples of how two cell masks in consecutive frames (previous, next) with little to no overlap become heavily overlapped after interpolation during **(H)** lateral rotation **(I)** rotation and cell growth **(J)** linear displacement. Yellow arrow = displacement direction. **(K)** spore release, where interpolation increases overlap between the previously sporulated mother cell mask and its released spores. **(L)** Representative comparison of single cell tracking without (top) and with (bottom) frame interpolation during exponential proliferation after germination. Yellow arrows = alternative tracking. Orange arrows = tracking error due to sudden cell displacement **(M)** Visualization of representative single cell tracks during exponential proliferation as solid masks for birth (blue), final location (pink), and mask contours for all intermediate masks (light blue). **(N)** Single cell velocities (µm / min) calculated using the locations of cell centroids after FIEST tracking of exponential growing yeast microcolonies.

We concluded that life cycle-stage segmentation enabled the detection of morphologically diverse cellular states in *S. cerevisiae*’s sexual life cycle.

### Frame Interpolation Enhanced Single-cell Tracking (FIEST) during life cycle transitions

To track single yeast cells during life cycle transitions we developed <u>F</u>rame Interpolation <u>E</u>nhanced <u>S</u>ingle-cell <u>T</u>racking, FIEST, an algorithm that (1) uses the overlap of masks between consecutive images to track cells and (2) ensures the overlap of cell masks even if no overlap exists between consecutive images. The latter was achieved by generating synthetic images through the interpolation of consecutive images using <u>R</u>eal-time Intermediate <u>F</u>low <u>E</u>stimation (RIFE ^62^). The goal of the interpolated synthetic images is to recreate not-recorded intermediate cellular states that occurred between the recorded frames (**Fig 3 A**). The segmentation of interpolated synthetic images can provide additional overlapping cell masks to ease cell tracking. Similar approaches have been developed to track objects in low frame rate videos ^78, 79^, enhance computer animation ^80, 81^, or improve temporal resolution in microscopy ^82^. By focusing on the overlap of single cell masks, FIEST avoided tracking requirements that can change during life cycle transitions, such as overreliance on cell centroids ^55^, matching of cell structural information features ^83^, or particle linking calculations ^84^.

To assess whether the synthetic images generated accurate segmentations for tracking purposes, we down sampled the frames of a full life cycle time series, generated the missing frames through interpolation and compared the segmentation results derived from synthetic images against the real images (**Fig 3 B**). The resulting interpolated images recapitulated real images (**Fig 3 C**, Video S3) as judged by structural similarity values (SSIM) > 0.95 throughout life cycle transitions **(Fig S3 A, B).** Synthetic and real images produced segmentations with similar single pixel cell probabilities (**Fig 3 D**) and similar single cell masks (**Fig 3 E**). Masks derived from interpolated images had > 0.9 AP when compared to masks derived from real images (**Fig 3 F**) independently of yeast strain background **(Fig S3 C)**. Hybrid time series containing real and synthetic interpolated images depicted the potential intermediate locations of cells during abrupt life cycle transitions such as exponential growth during germination (**Video S4**). The segmentation of hybrid time series containing real and synthetic interpolated images increased the overlap of cell masks across abrupt cell movements such as during irregularly-shaped mating events (**Fig 3 G**), lateral rotations (**Fig 3 H, Fig S3 D**), rotations combined with cell growth (**Fig 3 I, Fig S3 E**), linear displacement (**Fig 3 J, Fig S3 F**), and also help to increase the overlap between released spores and their sporulated mother cell (**Fig 3 K, Fig S3 G**).

To assess whether single cell tracking was improved by including segmentations from synthetic interpolated images, we scored tracking success before and after interpolation during spore germination **(Video S5)**; or during exponential growth in rich medium where cells simultaneously grow and drift while budding **(Video S6)**. FIEST improved single cell tracking success from 74 ± 5 %, to 97± 3 % (n = 680 tracked cells) (**Fig 3 L**), even for cells that were transported through irregular trajectories over distances larger than ten times their size (**Fig 3 M, Fig S3 H,)**, and with velocities of up to 20 µm / min, as measured by the euclidean distance between the coordinates of the cell’s birth and final location (**Fig 3 N**).

We concluded that FIEST enable the tracking of single cells through morphological transformations and abrupt displacements using minimal hyperparameter tuning.

### Cell growth precedes meiosis and exit from non-proliferative states in *S. cerevisiae*’s life cycle

Using life cycle-specific segmentation models and FIEST, we tested whether tracking *S. cerevisiae*’s life cycle under microfluidics conditions recapitulated morphology and size patterns previously reported for individual life cycle transitions ^85, 86^ ^51, 85–88^. F0 diploids (ancestors) were tracked individually through sporulation and up to ascus breakage after which the F1 diploids (descendants) were tracked independently (**Fig 4 A, Video S7).**

**Fig 4.**
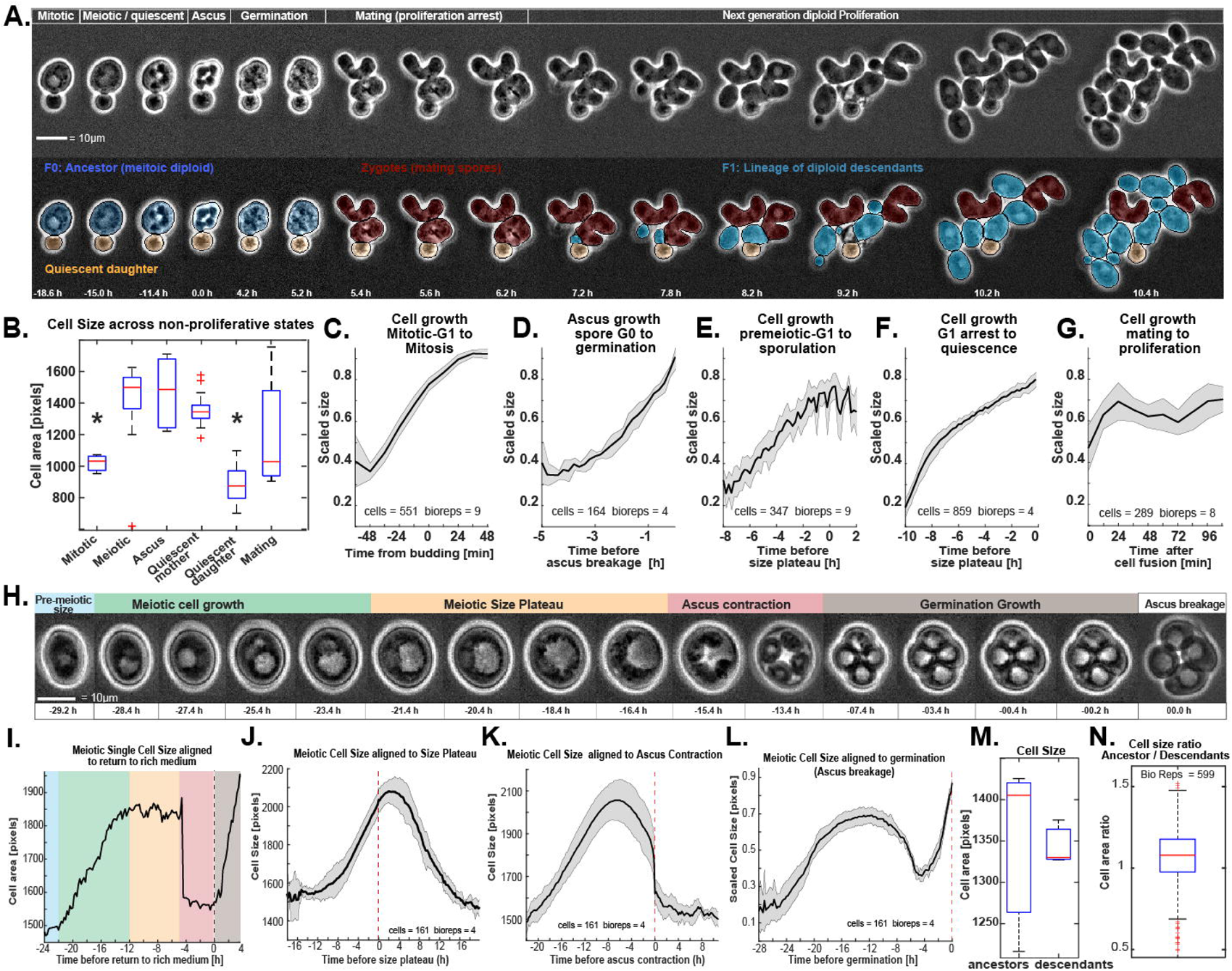
Cell growth regulation in *S. cerevisiae*’s sexual life cycle. **(A)** Concatenated time series micrographs with computationally aligned yeast cells during sexual life cycle. (top) phase images; (bottom) tracked single cell masks overlay with cell outlines depicted as solid black contours on phase image **(B)** Boxplot comparison of cell size at across non-proliferative life cycle stages. *Mitotic* = onset of budding; *meiotic* = ∼ 6 hours before spore formation; *ascus* = four-spore asci, *quiescent mother* = cells that entered quiescence after 10 hours exposure to starvation; *quiescent daughter* = cells born after stress after 10 hours. Mating = size at cell fusion. **(C)** Average cell size around the time point of budding in rich medium **(D)** Average cell size time series of tetrads after exposure to rich medium aligned to the timepoint of ascus breakage (germination) **(E)** Average cell size time series of meiotic cells aligned to the timepoint of size plateau during meiosis **(F)** Average cell size time series of quiescent mother cells aligned to the timepoint of size plateau during quiescence **(G)** Average cell size time series of zygotes (fusing mating cells) aligned to the timepoint of cell fusion during mating. **(H)** Concatenated time series micrographs with a computationally aligned representative yeast cell undergoing size changes during meiosis, sporulation, and germination **(I)** Representative single cell size during sporulation and germination. Background color matches top labels in H. **(J)** Average size of meiotic cells during starvation and sporulation aligned to the onset of the meiotic size plateau **(K)** Average size of meiotic cells during starvation and sporulation aligned to ascus contraction **(L)** Average size of meiotic cells during starvation, sporulation, and return to rich medium aligned to germination **(M)** Comparison of average cell size at the population level between ancestor and descendants cells **(M)** Average ancestor / descendant mitotic single cell size ratios. Solid lines with shaded area = average ± 95% confidence intervals. Asterisk = P < 0.05, K-S test, n > 3. Box plots display data from biological replicates, n > 3: central mark, median; box bottom and top limit, 25th and 75th percentiles; whiskers, most extreme non-outlier values, read crosses = outliers. Asterisks = p > 0.05, KS test, n >3.

Comparison of diploid cell size across *S. cerevisiae*’s life cycle confirmed previous reports of mitotic cells at budding onset being smaller than meiotic cells in pre-meiotic G1 ^35^ and quiescent mother cells; in addition, we found tetrads to be larger than mitotic cells, quiescent daughter cells (cells born under starvation) to present the smallest diploid sizes, and mating cells to have largest size variation (p<0.05, KS test, n>4) (**Fig 4 B**). Cells showed significant growth during life cycle transitions out of G1/G0 states, such as during the transitions from mitotic G1 into mitosis (**Fig 4 C**), from the G0 spore state into germination (**Fig 4 D**), from premeiotic-G1 into meiosis (**Fig 4 E**), and also from G1 arrest into quiescence (**Fig 4 F**) but not during the transition from mating into mitosis (**Fig 4 G**). Under starvation conditions, meiotic, and quiescent cells differed in several morphological and optical features at specific periods (Fig S4 A-C), and all cells became more spherical (Fig S4 D).

Single cell tracking revealed a characteristic oscillation in cell size during sexual differentiation (**Fig 4 H**, Fig S4 E), which consisted of an initial lag phase after starvation onset, followed by cell growth, a transient meiotic size plateau, and an abrupt size collapse due to ascus contraction that was reversed by spore growth in the ascus after transfer to rich medium (**Fig 4 I**, Fig S4 F). Meiotic cell size increased 28 ± 7 % (n=5) above mitotic levels as judged by alignment to the onset of the size plateau (**Fig 4 J**). Alignment to the point of size collapse revealed that an abrupt ascus contraction during spore maturation reset size to pre-meiotic levels (**Fig 5 K**). Alignment of meiotic cells to the germination time point showed that the ascus grew up to 33 ± 10 % before ascus breakage and spore release (**Fig 5 L**). The meiotic-germination size oscillation was consistent with alternative size measurements (Fig S4 G), and was different from oscillations in optical cell features (Fig S4 H).

**Fig 5.**
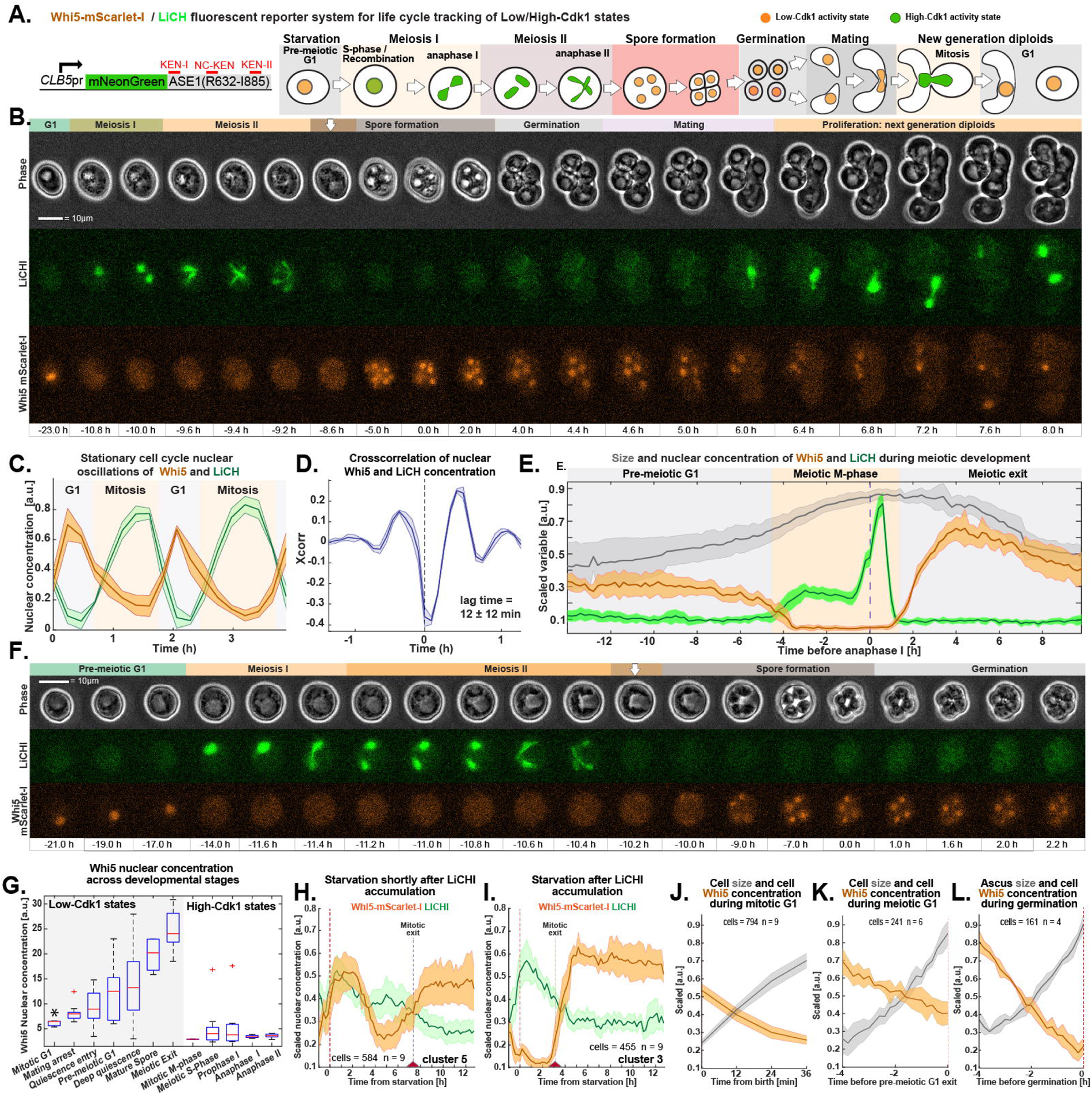
Continuous tracking of Cdk1-states across sexual life cycle stages. **(A)** Schematic of a fluorescent sensor system to track low- and high-Cdk1 states across life cycle stages. Solid circles = nuclei. NC-KEN = Non-canonical KEN destruction motif; green = LiCHI (*CLB5pr-NLS-mNeonGreen-ASE1(R632-I885));* orange = Whi5-mScarlet-I (Whi5-mSC). **(B)** Concatenated time series micrographs with a computationally aligned representative yeast life cycle. Top, phase images; middle, LiCHI fluorescence micrographs; bottom, Whi5-mSC fluorescence micrographs **(C)** Average nuclear concentration of Whi5-mSC (orange) and LiCHI (Green) during unrestricted proliferation in rich medium in aligned cell time series **(D)** Average crosscorrelation of Whi5-mSC and LiCHI nuclear concentration during proliferation. **(E)** Scaled size (gray) or Whi5-mSC (orange) and LiCHI (green) nuclear concentration during the transition from proliferation arrest in response to stress (pre-meiotic G1) into meiosis (Meiotic M-phase) and sporulation (Meiotic exit) **(F)** Concatenated time series micrographs with a computationally aligned representative yeast life cycle depicting (top) phase images, (middle) meiotic nuclear divisions indicated by nuclear LiCHI or non-proliferative states indicated by nuclear Whi5-mSC **(G)** Boxplot comparison of Whi5-mSC nuclear concentration across life cycle stages identified using LiCHI/Whi5 fluorescent sensor system **(H-I)** Average Whi5-mSC (orange) and LiCHI (Green) nuclear concentration during the transition from proliferation into starvation (dotted line) in *k*-means clustered cells according to nuclear Whi5 concentration. Cells experienced starvation **(H)** shortly before accumulating LiCHI (entry into mitotic M-phase) or **(I)** after LiCHI accumulation (M-phase). **(J-L)** Scaled average nuclear Whi5-mSC (orange) concentration and size (gray) during exit from non-proliferative states aligned to **(J)** cell birth during exit from G1, **(K)** pre-meiotic G1 exit, or **(L)** germination during the exit from the G0 spore state. Solid lines with shaded area = average ± 95% confidence intervals. Box plots display data from biological replicates: central mark, median; box bottom and top limit, 25th and 75th percentiles; whiskers, most extreme nonoutlier values. Asterisks = p > 0.05, KS test, n >3.

Average size and morphological features were statistically indistinguishable between ancestor and descendant diploid cells at the population level (**Fig 5 M**, Fig S4 I). To assess transgenerational differences between ancestor and descendants at the single cell level, we calculated the ratios for morphological properties between each ancestor cell and its descendants. Most ancestor/descendant ratios were close to unity and fell within the 95 % confidence interval of the cell lineage distribution, indicating that descendants were not significantly different from their ancestor cell in SK1 (**Fig 4 N**, Fig S4 J) or W303 (Fig S4 K) strain backgrounds. However, outlier descendants were identified in almost every lineage, revealing that in the next diploid generation single cells had a probability ≤ 4 ± 7 % (n=599) to be significantly different from its ancestor’s morphology.

We concluded that our single cell tracking system recapitulated previously measured cell size patterns during specific life cycle transitions, and, in addition, showed that *S. cerevisiae*‘s sexual life cycle comprises periods of cell growth during transitions out of non-proliferative states. Cell size and morphology persisted across generations at the population level, but morphological outliers were constantly present at the single cell lineage.

### Cdk1-States and Whi5 levels are differentially regulated across life cycle stages

To assess the coordination of cell growth with cell division during *S. cerevisiae*’s life cycle transitions, we developed a fluorescent sensor system to track high-and low-Cdk1 activity states across all life cycle stages. To visualize Low-Cdk1 activity states (mating, quiescence, mitotic G1, premeiotic-G1, and the spore state) ^31^, we tracked the nuclear localization of a fluorescently-tagged Whi5 transcriptional repressor. To visualize High-Cdk1 activity states (DNA replication, mitosis, meiotic recombination, meiosis I, meiosis II), we generated a fluorescent sensor for both mitotic and meiotic cell division, *CLB5pr-mNeonGreen-Ase1(R632-I885)* or <u>L</u>ife <u>C</u>ycle <u>H</u>igh-Cdk1 Indicator (LiCHI). LiCHI was composed of a mNeonGreen fluorophore expressed from the mitotically and meiotically-regulated *CLB5* promoter and fused to a C-terminal fragment of the protein ASE1, which provides nuclear targeting ^89^ and contains three degradation sequences, KEN motifs, recognized by the ubiquitin ligase APC/C-Cdh1 ^44^ (**Fig 5 A**). The nuclear occupancy of Whi5 and LiCHI was anticorrelated across *S. cerevisiae’s* life cycle stages (Fig 5 B**, Fig S5 A, Video S8**). During mitotic proliferation Whi5 and LiCHI generated stationary time series (Fig 6 C**, Fig S5 B)** with significant negative cross-correlation and a lag time of 12 ± 12 min as cells oscillated between Low-Cdk1 activity (G1 phase) and High-Cdk1 activity states (S/G2/M phases) (**Fig 5 D, Fig S5 C**). According to the nuclear Whi5 recruitment pattern, and other cell cycle markers (**Fig S5 D,E**), LiCHI accumulated during G1 exit and was destroyed during mitotic exit, recapitulating the mitotic expression profile of the CLB5 gene ^90, 91^ and the anaphase destruction of the Ase1 protein ^92^. We operationally defined mitotic G1 exit as the point of initial LiCHI accumulation and mitotic M-phase exit as the time point of full LiCHI destruction (**Fig S5 B**). Alanine substitutions in the KEN destruction motifs of the LiCHI reporter were sufficient for its stabilization during cell division, confirming that its degradation is APC/C-Cdh1-dependent degradation (**Fig S5 D,E**).

**Fig 6.**
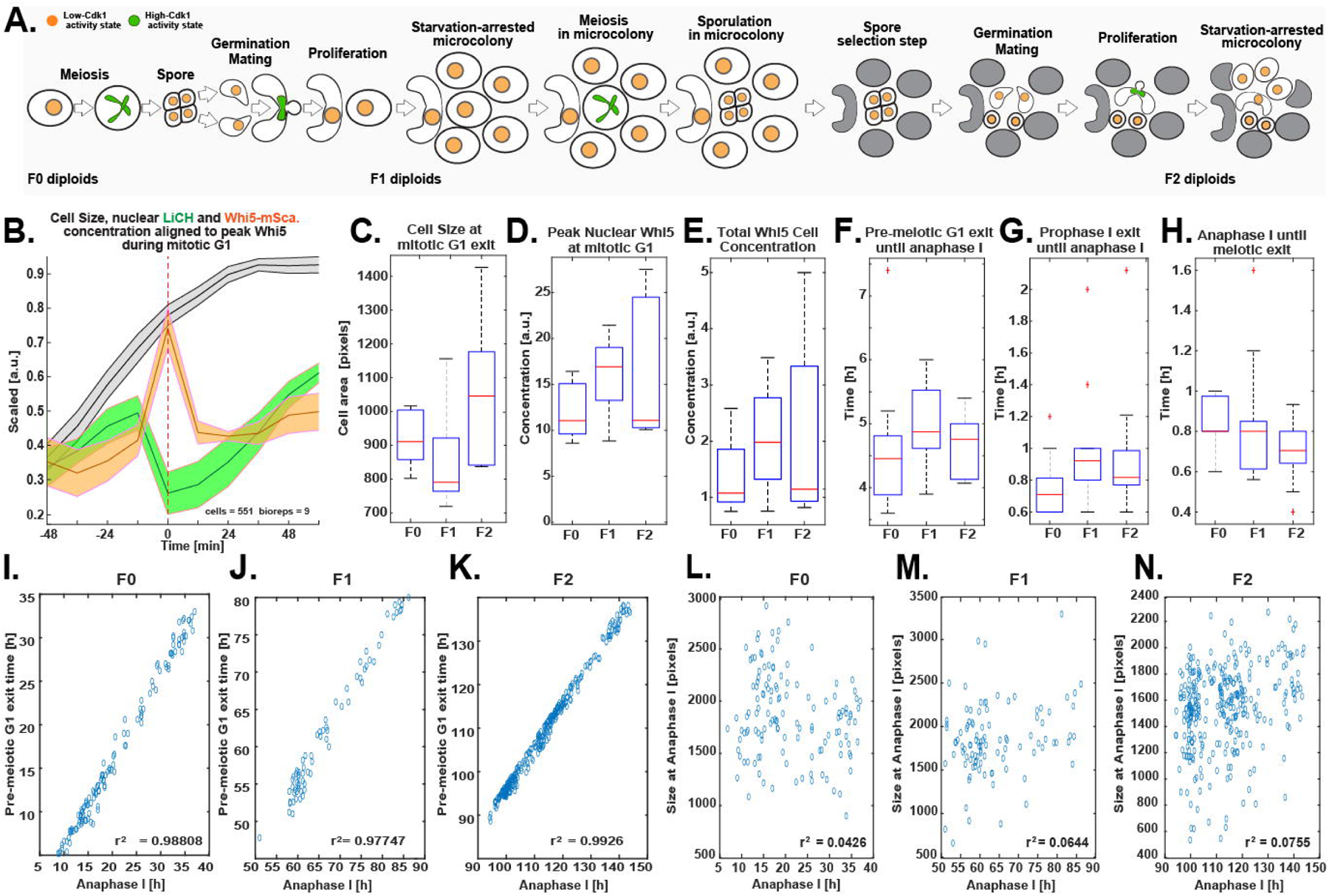
Continuous imaging of cell growth and cell division across sexually-reproducing yeast generations. **(A)** Schematic microfluidic induction of sequential *S. cerevisiae* sexual life cycles. Solid circles = nuclei; green = LiCH, orange = Whi5-mSC. **(B)** Representative scaled average size, Whi5-mSC or LiCHI nuclear concentration during the transition from cell birth into the first mitotic division aligned to the peak of Whi5 concentration during G1 **(C-E)** Boxplot comparison of the first mitotic division in three sexually reproducing generations (F0-F2) in terms of **(C)** cell size, **(D)** peak nuclear Whi5 levels, **(E)** total Whi5 cell concentration. **(F-H)** Boxplot comparison of the kinetics of meiotic divisions in three sexually reproducing generations (F0-F2) in terms of the duration between the time point of anaphase I and **(F)** pre-meiotic G1 exit, **(G)** prophase I exit, **(H)** meiotic exit. **(I-K)** Correlation between the time of anaphase I and pre-meiotic G1 exit in **(I)** F0, **(J)** F1, and **(K)** F2 generations. **(L-N)** Correlation between the time of anaphase I and cell size at anaphase I **(L)** F0, **(M)** F1, and **(N)** F2 generations.

During meiosis, single cell time series showed that Whi5 was excluded from the nucleus before and during meiotic divisions; whereas LiCHI was expressed in two waves that recapitulated the bimodal meiotic upregulation of the CLB5 gene at meiotic G1 exit and at prophase I exit ^93^ (**Fig 6 E,F**). We operationally defined pre-meiotic G1 exit as the point of initial LiCHI accumulation, the exit from prophase I as the onset of the second LiCHI accumulation peak, anaphase I as the first irreversible separation of the LiCHI nuclear signal, anaphase II as the second irreversible separation of the LiCHI nuclear signal, and meiotic exit as the time point of peak Whi5 nuclear accumulation after meiotic nuclear divisions **(Fig S5 F)**.

LiCHI was not significantly destroyed between meiosis I and meiosis II, enabling the tracking of both meiotic nuclear divisions (**Fig 5 F, Fig S5 G**), and supporting previous bulk assays reports ^95, 96^ indicating that APC/C-Cdh1 or its meiosis-specific counterpart APC/C-Ama1 are not active between meiosis I and II^94, 95^. Consistent with this view, nuclear Whi5 was not detected between meiosis I and meiosis II, regardless of Whi5 fluorescent tags or *in-silico* alignments (**Fig S5 G-I**), indicating an uninterrupted High-Cdk1 kinase state from pre-meiotic G1 exit until meiotic exit.

Comparison of Whi5 levels across life cycle stages revealed a strong developmental regulation of nuclear Whi5 in Low-Cdk1 states. Mitotic G1 presented the lowest nuclear Whi5 concentration, while meiotic exit and the spore state had the highest accumulation of nuclear Whi5 (KS-test, p < 0.05, n=7, **Fig 5 G**). Cells in deep quiescence (after 10 hours in starvation) had the most variable nuclear Whi5 levels. During high-Cdk1 states, including anaphase I and anaphase II, mitotic M-phase, meiotic S-phase, and prophase I, nuclear Whi5 levels were low and not significantly different (p > 0.05, n=7, **Fig 6 G**). The total concentration of Whi5 per cell, however, differed significantly between mitotic and non-mitotic states ( p < 0.05, n=7, **Fig S5 J**).

The only period when both nuclear Whi5 and LiCHI were undetectable was briefly after anaphase II during meiotic exit (**Fig 5 B, F**, top white arrow, **Fig S5 K**). Conversely, both Whi5 and LiCHI could be simultaneously present in the nucleus of cells that faced starvation during the last mitotic M-phase before meiosis entry (**Fig S5 L**). Whi5 time series clustering of proliferating cells exposed to starvation before meiosis confirmed that nuclear Whi5 levels increased in cells that faced starvation shortly after accumulating LiCHI (G1 exit), in agreement with previous reports ^48^ (**Fig 5 H, Fig S5 M**). By contrast, cells that faced starvation later in mitosis, after LiCHI accumulation, maintained Whi5 / LiCHI cell cycle coordination (**Fig 5 I, Fig S5 M**).

Total Whi5 concentration decreased during transitions from non-proliferative states such as mitotic G1 exit, pre-meiotic G1 exit, and the G0 spore state exit during germination, as measured by plotting the total Whi5 cell concentration aligned to the time of cell birth (**Fig 5 J**), pre-meiotic G1 exit (**Fig 5 K**) or germination (**Fig 5 L**), consistent with a model for Whi5 dilution during transitions out of non-proliferative states ^49^. By contrast, the total concentration of LiCHI levels did not decrease during transitions out of non-proliferative states (**Fig S5 N**).

We concluded that *S. cerevisiae’s* life cycle was characterized by developmentally regulated nuclear Whi5 concentration during non-proliferative states, a continuous High-Cdk1 state during meiotic divisions, and transitions between Low- and High-Cdk1 states marked by a decrease in the total Whi5 concentration.

### Coordination of cell division and growth persists across sexually-reproducing yeast generations

To assess the coordination of cell growth and cell division across sexually reproducing yeast generations, we induced three sequential sexual life cycles and tracked ancestor (F0, parental) and descendant diploid cells (F1-F2, filial) bearing the LiCHI/Whi5 reporter system (**Fig 6 A, Video S9)**.

To compare size and Whi5 levels during mitotic cell division across generations at the population level, we aligned all cells in each diploid generation to the time point of peak nuclear Whi5 levels during G1 exit (**Fig 6 B).** Cell size (**Fig 6 C)**, peak nuclear Whi5 concentration (**Fig 6 D),** and total Whi5 concentration (**Fig 6 E)** remained statistically indistinguishable across generations (KS-test, p > 0.05, n > 4).

To measure the kinetics of meiotic cell division and meiotic cell size regulation across generations, we measured the intervals between anaphase I, pre-meiotic G1 exit, prophase I exit, and meiotic exit. The temporal coordination of meiotic events in ancestors and descendants was also indistinguishable (KS-test, p > 0.05, n > 4), as measured by the duration between anaphase I and pre-meiotic G1 exit (**Fig 6 F)**, prophase I exit (**Fig 6 G)**, meiotic exit (**Fig 6 H)** at the population level. Analysis of the correlation between the time of anaphase I and pre-meiotic G1 exit (**Fig 6 I-K)** or all other measured meiotic stages (**Fig S6 A-C**), confirmed a tight regulation in the timing of meiotic events in every generation, with coefficients of correlation > 0.99. In sharp contrast, the size of meiotic cells at each specific stage was not correlated to the timing of any meiotic event (**Fig 6 L-N**). Average cell size, nuclear and total Whi5 at pre-meiotic G1 exit, prophase I exit, and meiotic exit showed no significant difference across generations (KS-test, p > 0.05, n > 4) **(Fig S6 D-F)**.

We concluded that despite the dramatic changes in cell growth and Whi5 levels during *S. cerevisiae*’s sexual life cycle, the tight temporal coordination of cell growth and cell division persisted, under our conditions, through three generations of sexual reproduction.

## Discussion

This work combined microfluidics, frame interpolation-driven single cell tracking, and fluorescent sensors to quantitatively analyze single cells through sexually reproducing generations in a model eukaryote, *S cerevisiae*.

In our experimental set up two variations of the homothallic life cycle were distinguished, which have implications for allele homozygosity after sexual reproduction ^96, 97^. In the homothallic life cycle type A, germinating spores immediately mated inside their ascus, which forces sister spores to potentially share genetic combinations derived from meiotic recombination and promotes heterozygous diploid descendants^30^. In the homothallic life cycle type B, germinating spores underwent haploid cell divisions before triggering sister-sister and mother-sister mating, which forces mating between genetically identical cells and promotes homozygous diploid descendants. Therefore, the homothallic life cycle A promotes heterozygosity, whereas the homothallic life cycle B promotes homozygosity. Whether the homothallic life cycle of yeast can be directed towards A or B in response to environmental or nutritional conditions, remains to be tested.

FIEST showed that the life cycle of sexually reproducing microorganisms can be tracked despite drastic morphological transformations using pixel flow vector-driven algorithms ^61, 62, 71^. Life cycle stage-specific segmentation models in combination with deep learning frame interpolation could be used to track filamentous organisms, shape-shifting cell lines, or bacteria in complex microscopy time series. Although deep learning single cell tracking does not offer a refined mathematical model to represent cell movement of morphological transformations, FIEST could be potentially combined with bio-realistic generative AI image and video processing tools ^98–100^ to leverage data to formalize mathematical models for single cell behavior.

Cell growth was a constant during exit from non-proliferative states such as mitotic G1, pre-meiotic G1, and the G0 spore state, which is consistent with the Whi5/Rb dilution model that has been shown as a mechanism for size control in single-celled organisms and mammals ^101^. Our data suggests that Whi5 dilution through cell growth could be a general feature of the exit from dormant and non-proliferative states. Size increase has been observed during the exit from dormant states in organisms as diverse as quiescent stem cells and fungal plant pathogens ^102, 103^.

Nuclear Whi5 levels were life cycle stage-specific, which supports the existence of still not fully characterized stress and developmental-specific mechanisms to control nuclear Whi5 ^48, 104^. Different developmental states might set specific nuclear Whi5 levels in the same manner that DNA replication and chromosome segregation requires different levels of Cdk1 activity ^105^.

The negatively crosscorrelated mitotic oscillations of Whi5/LiCHI fluorescent reporter system supports the standard mitosis model as transitions between mutually exclusive Cdk1 states ^22, 106, 107^. In meiosis, however, the Whi5/LiCHI system failed to detect the accumulation of nuclear Whi5 or the destruction of LiCHI between meiosis I and meiosis II, indicating that the transient activation of Cdk1 counteracting mechanisms during anaphase I, such as the activation of the phosphatase Cdc14 and APC/C-Cdc20 ^95^, only represent a perturbation in a steady state of High-Cdk1 activity. Consistent with this interpretation, recent phosphoproteomics studies comparing mitotic exit to anaphase I, found that Cdk1 targets largely remained phosphorylated at the end of meiosis I ^108^.

Sexual reproduction in yeast has been proposed to reset information accumulated from previous life cycles ^109, 110^; our results support this view, showing that when sexual life cycles were induced cyclically, each generation maintained high temporal coordination in cell division and growth regardless of cyclic nutritional stress, in contrast to asexual life cycles based on cellular quiescence or dormancy ^36^. To further explore these open questions, manufacturing of wider aspect ratio microfluidic devices will increase the number of consecutive sexual life cycles that can be recorded, opening the doors to quantitative live-cell studies on transgenerational biology and evolution by sexual reproduction.

## Materials and Methods

### Yeast strain and plasmids

Experiments were carried out with three different *Saccharomyces cerevisiae* strain backgrounds: ATCC WT SK1 (*MAT*a/α *HO gal2/ gal2 cup^S^/cup^S^ can1^R^ / can1^R^ BIO*), laboratory SK1 (*ho::LYS2 lys2 ade2Δ::hisG trp1Δ::hisG leu2Δ::hisG his3Δ::hisG ura3*), and W303 (*leu2-3,112 his3-11,15 ura3-1 trp1-1 can1-100 ade2-1*). Statistical comparisons were only performed between isogenic strains. All promoter and open reading frames used in plasmid construction were derived from *S. cerevisiae* W303 using high-fidelity PCR amplification, subcloned into pGEM-T, and confirmed by sequencing and restriction analyses. Tagged fluorescent proteins were homozygous unless otherwise stated. Diploids were created by mating appropriate haploids obtained by tetrad dissection or by transformation with PCR tagging cassettes, deletion cassettes, or plasmids, using the polyethylene glycol/lithium acetate protocol ^111^. One-step PCR was used for gene deletion of *SFP1* and for C-terminal fluorophore tagging *WHI5* with monomeric mScarlet-I or monomeric Kusabira Orange kappa, mKOκ, and C-terminal tagging of *CDC10* with monomeric Cyan Orange Fluorescent protein, mCyOFP1, using cassette plasmids described previously ^51^. *CLN3* and *VAC17* deletion strains have been described elsewhere ^51^. Genomic integrations and deletions were confirmed by PCR with appropriate checking primers (Table S2).

The *CLB5pr-mNeonGreen-Ase1(R632-I885)* LiCHI plasmid was created by ligating the following fragments: (1) a *Sac*I/*Pac*I–flanked CLB5 promoter sequence (-582 to −1); (2) a *Pac*I-*Asc*I–flanked mNeonGreen (yeast optimized) fluorophore from pYLB10 ^51^; (3) an *Asc*I-*Not*I–flanked C-terminus of ASE1 (R632-I885); and (4) a pRS305 backbone cut with *Not*I and *Sac*I.

The non-destructible version of the *Ase1(R632-I885)* fragment was produced by creating three pGEM plasmids containing PCR-mutagenized sequences of the APC/C-Cdh1 recognition sequences (KEN motifs). Primers to mutagenize the first KEN sequence changed lysine 642, glutamic acid 643, and asparagine 644 to alanine and produced a fragment flanked by *Pac*I-*Sac*II. Primers to mutagenize the second non-canonical Cdh1-recognition sequence changed arginine 760, phenylalanine 763, and asparagine 768 to alanine, according to ^92^, creating a *Sac*II-*Mfe*I flanked fragment. Primers to mutagenize the third KEN sequence changed lysine 798, glutamic acid 799, and asparagine 800 to alanine creating *Mfe*I-*Pst*I flanked fragment. The fragments were triple ligated to a *Sac*II and *Pst*I cut backbone from *MET3pr-mNeonGreen-Ase1(R632-I885),* which has been described elsewhere ^36^. The *MET3pr-mNeongGreen-CDH1* plasmid resulted from a pRS305 backbone triple ligated to a *Sac*I-*Pac*I flanked MET3-promoter (-517 to -1), a *PacI*-*AscI* mNeonGreen fragment from pYLB10 and a *PacI*-*NotI* flanked *CDH1* ORF including 104 bp after the stop codon. Linearization with *Age*I allowed integration of pRS305 plasmids at LEU2 locus.Table S1, S2, and S3 enumerate all yeast strains, primers, and plasmids, respectively.

### Growth conditions and microfluidics life cycle induction

For life cycle induction, yeast cells were inoculated at a starting concentration of OD_600_ = 0.03 in YPD (1 % yeast extract, 2 % peptone, 2% glucose), and grown overnight at 30 °C with orbital shaking at 295 rpm until reaching ∼ OD_600_ = 2; which corresponds to early stationary phase. The culture was spun down on a tabletop microfuge for 3 seconds to remove clumps, and 70 µl of the top layer were transferred to a microfluidics device with flow control (Y04C CellASIC plate with OniX controllers), pre-warmed at 25 °C. Meiotic induction was triggered by exposing the cells for 24+ hours to sporulation medium (SPO) at 0.6 psi / 8.3 kPa isobaric flow rate in the CellASIC device. SPO medium consists of 0.6% potassium acetate, 0.02 % sorbitol (from a 2 % w/v stock solution), 40 mg / L adenine, 40 mg / L uracil, 20 mg / L histidine, 20 mg / L leucine, 20 mg / L tryptophan, and adjusting the pH to 8.5 using a 0.25 M Na_2_HCO_3_ solution right before experiments. After sporulation, cells were exposed to a solution of 4 M LiCl containing 0.9 mg / mL Nystatin (from a 3 mg / mL DMSO stock solution) followed by 1 hour in sterile distilled water, which kills non-sporulated and quiescent cells. Spore germination was triggered using a flow rate of 0.3 psi / 4.2 kPa to switch cells to synthetic complete medium (SCD: 1 % succinic acid, 0.6 % sodium hydroxide, 0.5 % ammonium sulfate, 0.17 % yeast nitrogen base without amino acids or ammonium sulfate, 0.13 % amino acid dropout powder (complete), 2 % glucose) supplemented with 5 % YPD. This microfluidic protocol was sequentially repeated for experiments that image multiple sexual life cycles.

### Generating the training data set of morphologically diverse yeast shapes

Nocodazole M-phase synchronized cells were obtained from YPD cultures, 0.3 OD_600_, that were spun down and resuspended in 20 µg / ml nocodazole-containing SCD and cultured for 2 h until ∼ 90 % of cells were dumbbell-shaped. G1 synchronized cultures were obtained by centrifuging a YPD culture, OD_600_ =2, at 500 rpm for 2 min on a discontinuous PEG 5000:SCD-minus-dextrose three lawyered gradient (1:3, 1:27, 1:81) and taking the cells in the top 200 μl (∼ 90 % unbudded). To induce drastic morphology changes, cell bearing the *MET3pr-mNeongGreen-CDH1* construct were microfluidically cultured in SCD before transfer to SCD lacking methionine for 2 hours. For images depicted nutritionally stressed states, SK1, BY, and W303 cells we cultured in SCD for 2 hours before transfer to SCD medium lacking nitrogen sources or SCD containing 2 % glycerol or 2 % acetate or 2 % sorbitol as carbon source. Heterogenous cell size and shape was also induced by imaging W303 cells carrying deletions for *VAC17*, *SFP1*, and *CLN3* at variable cell densities. The training data set also contained images from diverse *S. cerevisiae* strains such ATCC BY (*MATa/MATα his3Δ1/his3Δ1 leu2Δ0/leu2Δ0 LYS2/lys2Δ0 met15Δ0/MET15 ura3Δ0/ura3Δ0*), and the industrial or environmentally-isolated strains YPS163 (Pennsylvania oak), CSH942 (Fleischmann’s baking yeast), UWOPS83.787.3 (Bahamas) and RM11 (Wine strain) ^112^, which were a gift from the Heil Laboratory (NCSU). Cellpose models ^60^ were trained on manually labeled sets of images containing cells in specific life cycle stages, focusing the models to detect only the pixel vector flows of specific life cycle stages to generate MatSeg and SpoSeg^60^. Labels for training the Budseg model were obtained from cells bearing a fluorescent septin ring component, Cdc10 ^77^. The fluorescent Cdc10 signal was used to automatically label budneck pixels in phase images **(Fig S2 B)**. Model training followed the steps described in ^113, 114^. The training was iterative until the models reach an average precision of at least 0.9 on the testing data set.

### Time-lapse microscopy

Microfluidics experiments were performed on an automated Zeiss Observer Z1 microscope controlled by ZEN pro software and with temperature control (Zeiss). Images were acquired at 12 min sampling rate using an 40X Zeiss EC Plan-Neofluar 40X 1.3 NA oil Ph 3 M27 immersion objective. Image focus was controlled using Definite Focus 3.0. Images were registered using an AxioCam 712 monochrome. An X-CITE XYLIS XT720S lamp (Excelitas Technologies) was used as light source. Tailored dichroic mirrors and bandpass filters ^51^ allowed sequential non-phototoxic fluorescence imaging using the following fluorophores and exposure times: the green-yellow fluorescent protein mNeonGreen ^115^ (100 ms), the mKusabira-Orange κappa fluorescent protein mKOκ ^116^ (200 ms), the synthetic-construct derived red fluorescent protein mScarlet-I mSC-I ^117^ (200 ms), and the long Stokes shift monomeric Cyan-Orange fluorescent protein mCyOFP1 ^118^ (35 ms), and phase-contrast for 20 ms.

### Image processing and phase or fluorescence quantification of cellular features

Images were collected using ZEN pro software (Zeiss) with 2 × 2 binning and exported in non-compressed 16-bit TIFF format, converted to double format before quantification, and median-filtered to exclude shot noise using *medfilt2()* with a 3 × 3 structuring element and symmetric padding option. After segmentation (see below), cell intensity was corrected for background fluorescence by subtracting the median value of the space not occupied by the cells in each image. A 2D Gaussian fit to the brightest pixel of nuclear proteins was used to calculate nuclear parameters ^119^. Nuclear concentrations were obtained using the custom function *Get_Sphere_Vol()*, which uses the equivalent diameter of a 2D mask to compute volume assuming a spherical shape. Cell morphological features such as extent, eccentricity, and equivalent diameter were obtained using the function *regionprops()* on cell masks. Refractivity-associated properties such as pixel correlation, homogeneity, and contrast were calculated on the phase images using the function *graycomatrix()* and *graycopropos()*. The septin ring was detected by measuring the standard deviation of Cdc10-mCyOFP1 fluorescence at the cell periphery as in ^36^. Code for fluorescence extraction is available at <u>MirandaLab/Parallelized_Fluorescence_Extraction: Parallelized_Fluorescence_Extraction</u> <u>(github.com).</u> All segmentations and interpolations were performed on a Dual Intel Xeon Silver 4216 (2.1GHz,3.2GHz Turbo, 16C,9.6GT/s 2UPI, 22MB Cache, HT (100W) equipped with an Nvidia Quadro RTX5000, 16GB, 4DP, VirtualLink (XX20T) outfitted with 64GB 4x16GB DDR4 2933MHz RDIMM ECC Memory and a 2TB primary NVMe SSD.

### Frame interpolation enhanced single-cell tracking (FIEST) on *S. cerevisiae*’s the life cycle

Each sequential pair of phase images from a time series was interpolated at least 16X using RIFE with default settings to create a hybrid time series of high-temporal resolution images ( equivalent to a sampling rate of < 1 min). Segmentations were performed on the hybrid time series composed of real and synthetic images using sequential detections by the custom pre-trained cellpose models: ProSeg, for generalist segmentation, MatSeg for segmentation of mating states after cell fusion, SpoSeg for spores and early germination stages, and BudSeg for detecting mother-daughter cell pairs based on their connection at the budneck. Full life cycle tracking involved the following steps on the hybrid image time series:

1. FIEST tracking: The SpoSeg, MatSeg and ProSeg segmentations of interpolated time series generated cell masks with unique indexes in each image. For every indexed cell mask at a given time point, the tracking algorithm projected the indexed cell mask from the previous image onto the cell masks in the next segmented image and identified the cell mask with the highest overlap. Once identified, the index of the cell mask in the next image is replaced by the index of the cell mask in the previous image, forcing each cell to acquire a unique index throughout the time series.
2. Correction of incomplete cell masks: ProSeg failures to correctly detect mating or sporulation events were identified as discrepancies in the number of cell masks or identified pixels by ProSeg, MatSeg or SpoSeg. Discrepancies between ProSeg, MatSeg, SpoSeg mask were resolved by using the cell mask that maintained the most coherent single cell track, for instance by preventing unnatural sudden drops or increases in cell size. This process often involved exchanging a faulty Proseg mask with a MatSeg or SpoSeg cell mask that achieved a better detection. After all potential corrections occurred, the tracked segmented images of the hybrid time series were downsampled ensuring that the final tracks only correspond to the real images. Final corrected single cell tracks could be a combination of Proseg and SpoSeg and MatSeg. Each cell detected by SpoSeg or MatSeg was assigned a unique identifier as sporulating or mating cell.
3. Mother-Daughter cell pair identification: the overlap between BudSeg masks and Proseg masks in tracked segmentations was used to find cell mask index pairs, if an index pair repeated more than three times in a tracked time series, the two masks were assigned a unique identifier as a mother and daughter pair.
4. Finding the indexes of the proliferative haploids that mated to form a zygote: each initial tracked cell mask for mating events, MatSeg, was skeletonized and overlapped with all previous time points in the ProSeg tracks, until finding the first time point where the area covered by the mating event contained two cell mask indices with an overlap ratio greater than 0.25 but less than 4. In case of ascus mating, this operation eventually leads to an overlap with SpoSeg masks, which indicates ascus mating.
5. Assign descendant cells to an ancestor sporulated cell: Descendants cells were determined by using a watershed algorithm that delineates an area around the centroids of germinating asci, allowing to affiliate ProSeg and MatSeg cell masks to a single germinating event. Source code available at: <u>MirandaLab/Full_Life_Cycle_tracking: Tracking for code for S. cerevisiae’s Life cycle (github.com)</u>.

### Time series analysis

To align cells to events that correspond to points of inflection on a time series, such as nuclear divisions or abrupt changes in size, the first derivative of the time series, denoised using *smooth(),* was calculated and the points of inflection were identified as changes in the sign of the derivative. Plateaus were identified as the first point of inflection before a region where the derivative sign stopped changing for at least three or more time points depending on the interval assessed.

The temporal coordination between stationary time series from different fluorophores was assessed by cross-correlation using the function *xcorr()*. Lag times were defined as the absolute highest xcorr value closest to zero. The function *kmeans()* created single cell time series clusters during starvation using the whole time series and the arguments: distance metric = correlation, replicates = 10, maximum iteration limit = 10000, and maximum number of clusters = 14. The ascus germination point was automatically marked as the time point when the first derivative of the ascus size time series reached its most negative value.

### Statistical analyses

Statistical analyses treated each independent microfluidic device as a biological replicate. Cell linages were considered a replicate if separated by more than 200 µm. Differences between populations of biological replicates were evaluated using the Kolmogorov-Smirnov tests *kstest2()*, with significance set at p < 0.05. Line plots correspond to the average of biological replicates surrounded by the 95 % confidence intervals represented as shaded area. Scatter plots depicted R-squared values calculated using *corrcoef()*. Box plots represented the median as the central mark, the 25^th^ and 75^th^ percentiles as the bottom and top limits of the box, and most extreme non-outlier values as whiskers. The function *isoutlier()* excluded outliers from statistical tests or identified outlier descendants in cell lineages.

### Computational alignment of cell micrographs and generation of videos

Phase contrast micrographs depicting single cells or single microcolonies were segmented using Proseg and their instance segmentation masks were turned semantic a segmentation mask (only cells or background), the centroid of the cell/microcolony binary mask was determined using *regionprops()* and the coordinates were used to crop a square around the cell mask and deposit the cell image in the middle of a numerical array composed of background pixels obtained from a randomized sample of the original image background (pixels not occupied by the cells). Images were automatically rotated so that the longest axis was Y matching the Y dimension of the last frame in the time series. Concatenation of such images produced the time series strips used throughout the paper. The pixels inside and around the cells were not modified by alignment and concatenation.

Videos were generated by scaling and normalizing the raw images across the time series data to remove noise and flickering, which facilitates focusing on single cells or their masks overlay. Single cell quantifications were never obtained from images processed for video.

## Code and Data availability

The original MATLAB image analysis code for single-cell tracking, quantifications derived from fluorescent extractions, time series visualization, generation of movies, computational alignment of cell micrographs, and automated statistical analyses is available at <u>MirandaLab (github.com)</u> including toy data sets to recapitulate results. The manually or algorithmically labeled datasets for general segmentation, mating, sporulation and budneck are available at NCSU libraries. The python code can be found in the repository <u>MirandaLab/yeastvision2: Deep-Learning enabled quantitative image analysis of for the yeast full life cycle</u> <u>(github.com)</u>. The*Yeastvision* GUI can be pip-installed from <u>yeastvision · PyPI</u> .

## Acknowledgements

In memoriam of Andreas Doncic. We thank Caiti Smukowski Heil for providing industrial and wild-type strains. We thank Monish Shah, Taevon Roach, Devika Venugopalan, Manav Patel, Aditi Koratpallikar, Sebastian Randomski, Aneka Goli, Isha Patania, and Bryn Merritt for data labeling and processing. This work was supported by grants R00GM135487 (OAM) and R01GM127614 (NEB) from the National Institute of General Medical Sciences of the National Institutes of Health, USA, and the Research Innovation Science Funds from NCSU-ORI/KIETS.

The authors declare no competing financial interests.

## Author contributions

S. Ramakanth: development of single-cell segmentation, interpolation, tracking, fluorescence extraction, data visualization, investigation, methodology, resources, validation, and statistical analyses. T. Kennedy: development of single-cell segmentation investigation, microfluidic assay development, methodology, resources, and validation. B. Yalcinkaya: complete GUI development, development of single-cell segmentation, methodology, resources, and validation. S. Neupane: microfluidic assay development, methodology, resources, and validation. N. Tadic: methodology, resources, statistical analyses, and validation. N. Buchler: conceptualization, software testing, supervision, funding acquisition, methodology, project administration, writing, and editing. O. Argüello-Miranda: conceptualization, data curation, formal analysis, funding acquisition, methodology, project administration, software development, validation, and writing.

---

**Fig S1.**
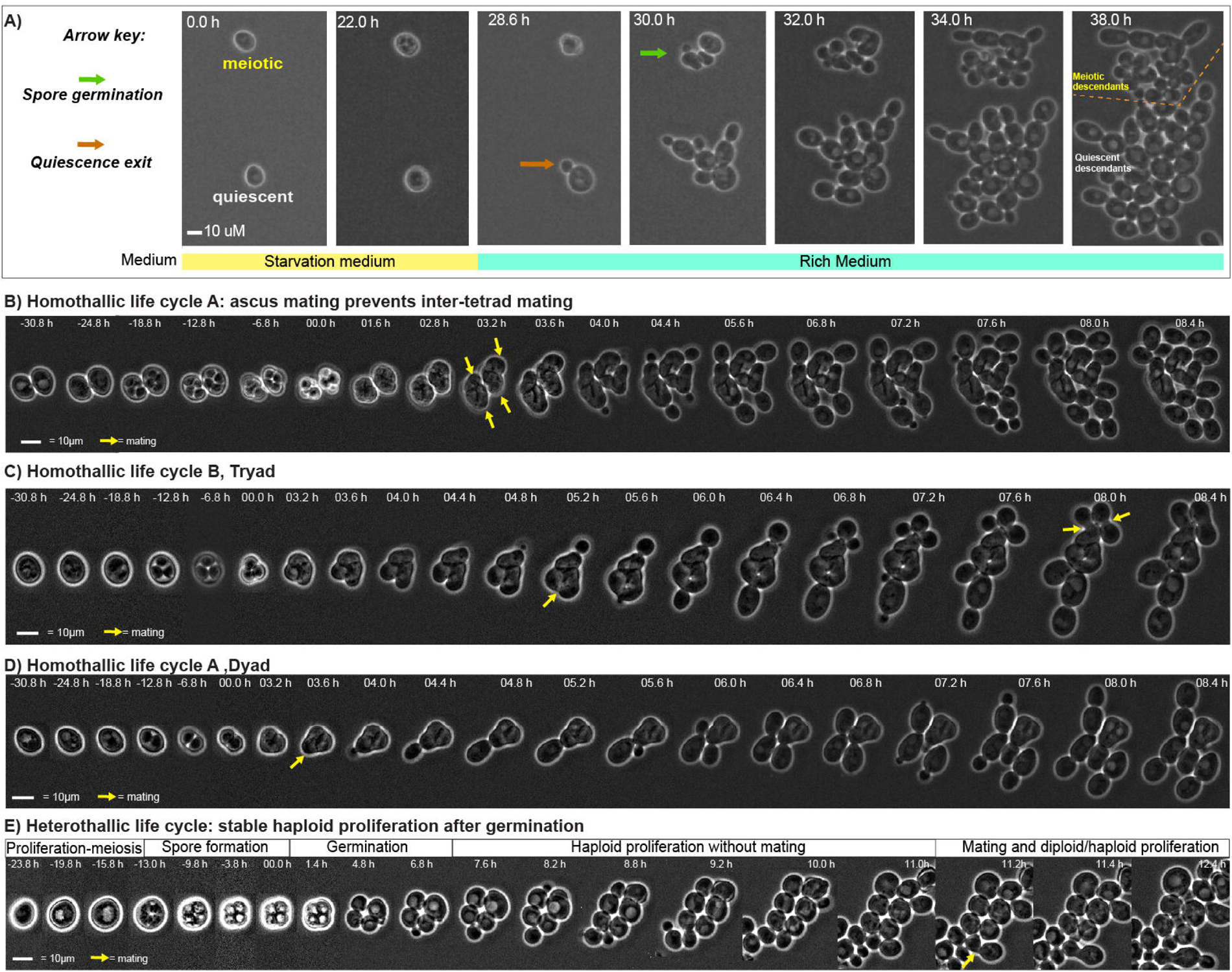
Microfluidics life cycle induction in the model eukaryote *Saccharomyces cerevisiae*. **(A)** Representative time series of a meiotic (top) or quiescent (bottom) cell during starvation and return to rich medium conditions. In the absence of the LiCl/nystatin treatment before return to rich medium, the descendants of the quiescent cells can outcompete the descendants of meiotic cells descendants and overpopulate the microfluidics device. Orange line = boundary between meiotic or quiescent descendants **(B-E)** Representative concatenated time lapse micrographs with computationally aligned yeast cells undergoing **(B)** homothallic life cycle type A, in which ascus mating prevented two sporulated cells in close proximity to undergo outbreeding. **(C)** Homothallic life cycle type B with three-spore asci (tryads), in which sporulation produced one diploid and one proliferative haploid cell that underwent proliferation followed by inbreeding. **(D)** Homothallic life cycle type A with two-spore asci (dyads), in which sporulation and germination directly leads to ascus mating and the production of a diploid without an intervening proliferative haploid phase. **(E)** Heterothallic life cycle in a laboratory W303 strain, which is unable to switch mating types due to an inactivating mutation in the HO endonuclease responsible for the recombination of MAT loci. The germinating spores produced four proliferative haploid cells that only engage in mating after several cell divisions.

**Fig S2.**
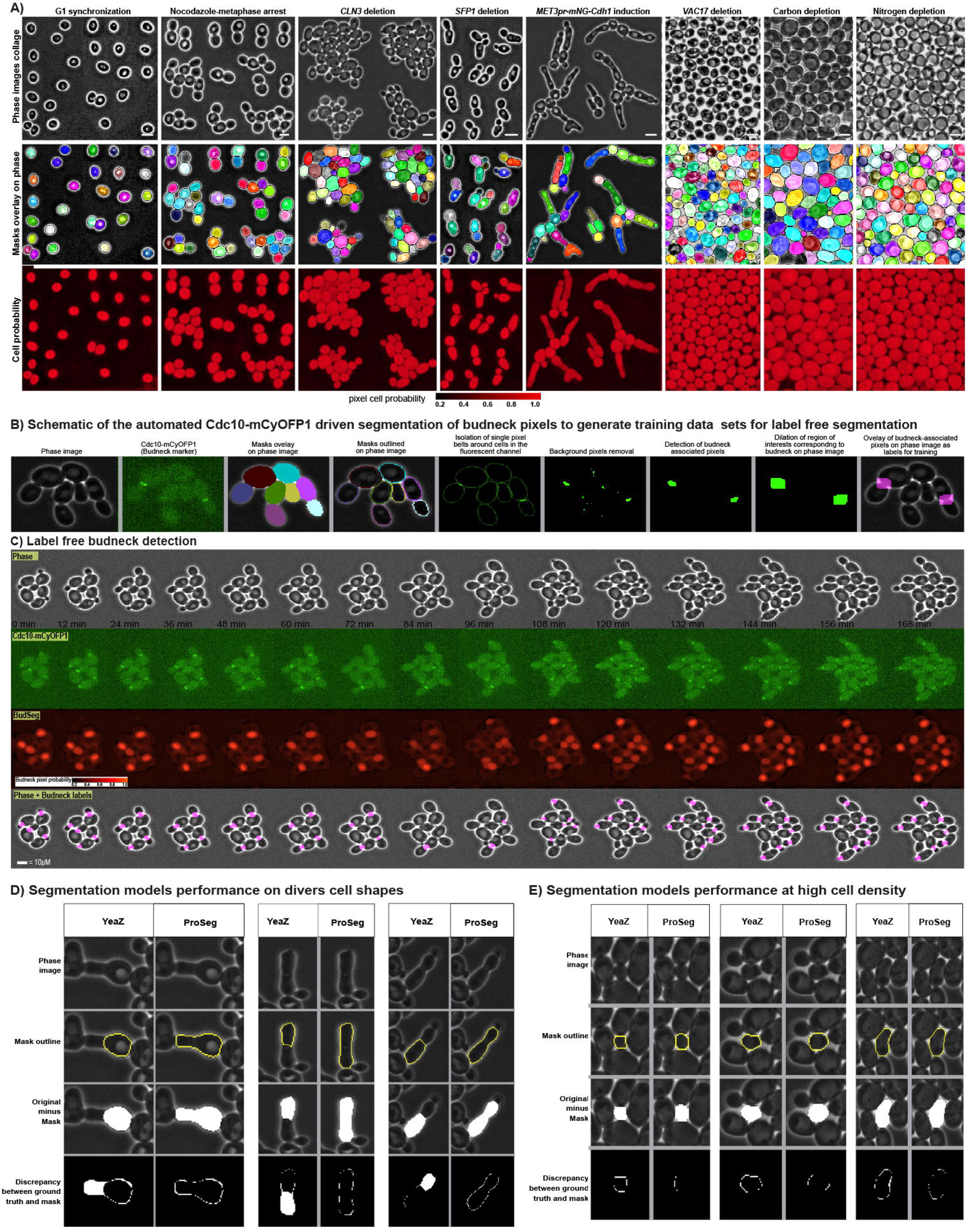
Life cycle stage-specific segmentation enables the detection of morphologically diverse stages in *S. cerevisiae*’s sexual life cycle. **(A)** Representative segmentations of yeast phenotypes that were included in the training data sets. First row, computationally aligned single cell collage. Second row, single cell masks overlayed on the phase image after segmentation. Third row, pixel cell probabilities generated as heatmap, which were used to generate the single cell masks above following the cellpose architecture. Size bar = 10 µM. Columns: *G1 Synchronization =* G1 cells obtained through gradient centrifugation; *Nocodazole-metaphase arrest* = cells after two hours exposure to 20 µM Nocodazole; *CLN3 deletion* = Enlarged cells due to the deletion of the size-controlling G1 cyclins *CLN3*; *SFP1 deletion* = small misshaped cells bearing a deletion of the ribosome biogenesis regulator *SFP1*; *MET3pr-mNeonGreen-Cdh1* = Cell shape perturbation by the conditional overexpression of the Hct1/Cdh1 APC/C activator; *VAC17 deletion* = organelle perturbation by deletion of vacuole/lysosome inheritance factor *VAC17*; *Carbon depletion* = Crowded cells after four hours under glucose depletion after transfer from rich SCD to SC-Acetate (SCA). *Nitrogen depletion* = Crowded cells after four hours under nitrogen source depletion after transfer from rich SCD to AM1 medium lacking nitrogen sources. **(B)** Schematic description of how the fluorescent septin ring marker Cdc10-mCyOFP1 was used to create a set of budneck-labelled phase images to train BudSeg, a label free segmentation model to detect the bud neck connection between mother and daughter cells from phase images. **(C)** Representative segmentation of: first row, a concatenated time series of micrographs with a computationally aligned yeast microcolony during proliferation; second row, fluorescent micrographs in the mCyOFP1 channel showing the septin ring marker, Cdc10-mCyOFP1; third row, BudSeg segmentations as budneck pixel probability heatmaps; fourth row, resulting budneck masks detected using only phase images. **(D-E)** representative visual comparisons of ProSeg and YeaZ segmentations emphasizing the discrepancies between the masks generated by each model and the human-generated ground truth segmentations for **(D)** elongated cell shapes and **(E)** in high density cultures.

**Fig S3.**
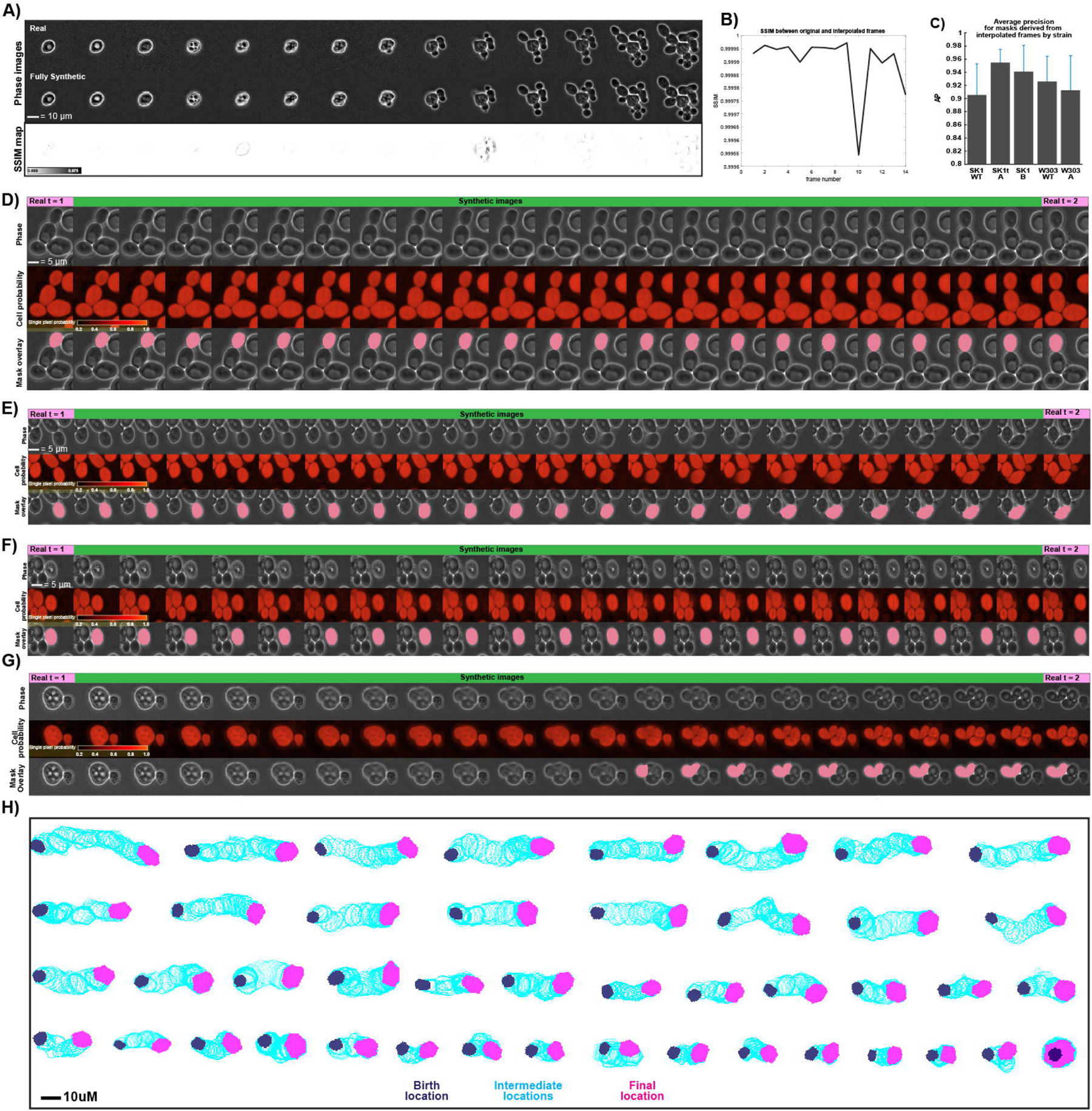
Single cell tracking across *S. cerevisiae*’s life cycle using Frame Interpolation Enhanced Single-cell Tracking (FIEST). **(A)** Real (top) and synthetic (middle) concatenated micrographs with computationally aligned yeast cells undergoing life cycle transitions and their (bottom) structural similarity index measurements (SSIM) as heatmaps. **(B)** Time series of the SSIM values obtained by comparing the original and synthetic frames in A. **(C)** Average precision bar plot comparing masks derived from real images to masks derived from synthetic images in multiple yeast strain backgrounds, n > 3. **(D-F)** Schematic comparison of two real consecutive frames (first and last) and their interpolated images during **(D)** lateral rotation **(E)** rotation and cell growth **(F)** linear displacement **(G)** spore release showing how interpolation eases the correct assignment of released spores to their sporulated mother cells. Top, phase images; middle, segmentation results as single pixel probability heatmaps; Bottom, tracked masks for the mating cells across real and interpolated images. **(H)** Visualization of diverse single cell tracks during exponential proliferation, with solid masks for birth (blue), final location (pink), and mask contours for all intermediate detections (light blue). Bar plots = mean ± standard deviation.

**Fig S4.**
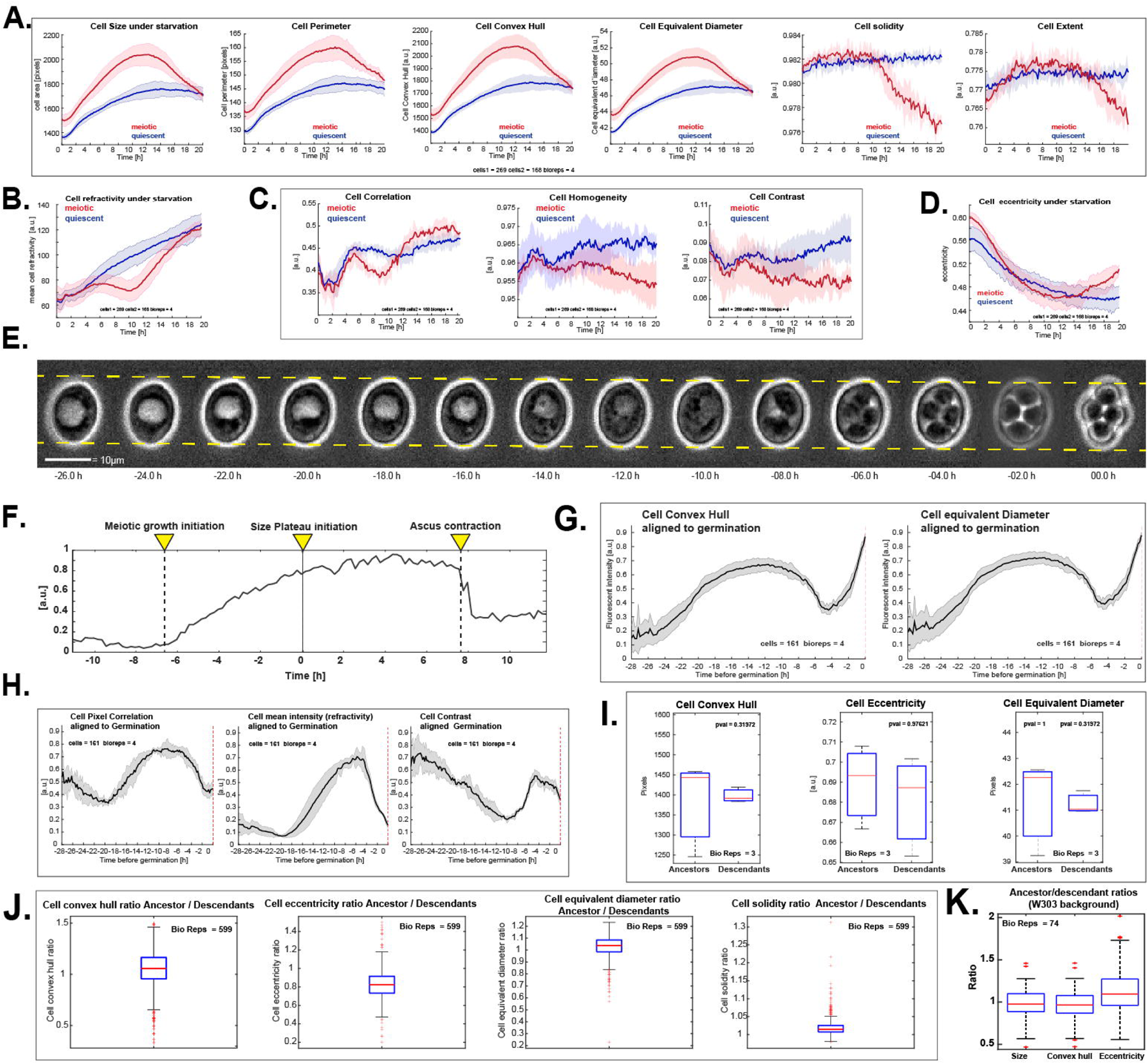
Cell growth regulation in *S. cerevisiae*’s sexual life cycle. **(A)** Average time series for morphological properties of meiotic and quiescent cells during starvation. From left to right: size, perimeter, convex hull, equivalent diameter, volume, solidity, extent. **(B)** Average refractivity of meiotic and quiescent cells during starvation. **(C)** Average time series for pixel features in meiotic and quiescent cells during starvation. From left to right: correlation, homogeneity, contrast. **(D)** Average eccentricity of meiotic and quiescent cells during starvation. **(E)** Concatenated time series micrographs with a computationally aligned representative W303 yeast cell during starvation and sporulation. Dotted lines = guides for size comparison **(F)** Representative single cell size during starvation and sporulation with indicated size landmarks **(G)** Average size of meiotic cells during starvation, sporulation, and germination expressed as the alternative size variables convex hull and equivalent diameter aligned to the timepoint of germination. **(H)** Average optical properties of meiotic cells during starvation, sporulation, and germination expressed as pixel correlation, cell mean intensity (refractivity), cell contrast aligned to the timepoint of germination. **(I)** Boxplot comparison of population level morphological properties such as convex hull, eccentricity, and equivalent diameter, between ancestor and descendant cells during mitosis **(J)** Boxplots of ancestor / descendant single cell ratios for morphological properties. From left to right: convex hull, eccentricity, equivalent diameter, solidity **(K)** Boxplots of ancestor / descendant single cell ratios in W303 strain background for morphological properties such as cell size, convex hull and eccentricity. Solid lines with shaded area = average ± 95% confidence intervals. Box plots display data from biological replicates: central mark, median; box bottom and top limit, 25th and 75th percentiles; whiskers, most extreme nonoutlier values.

**Fig S5.**
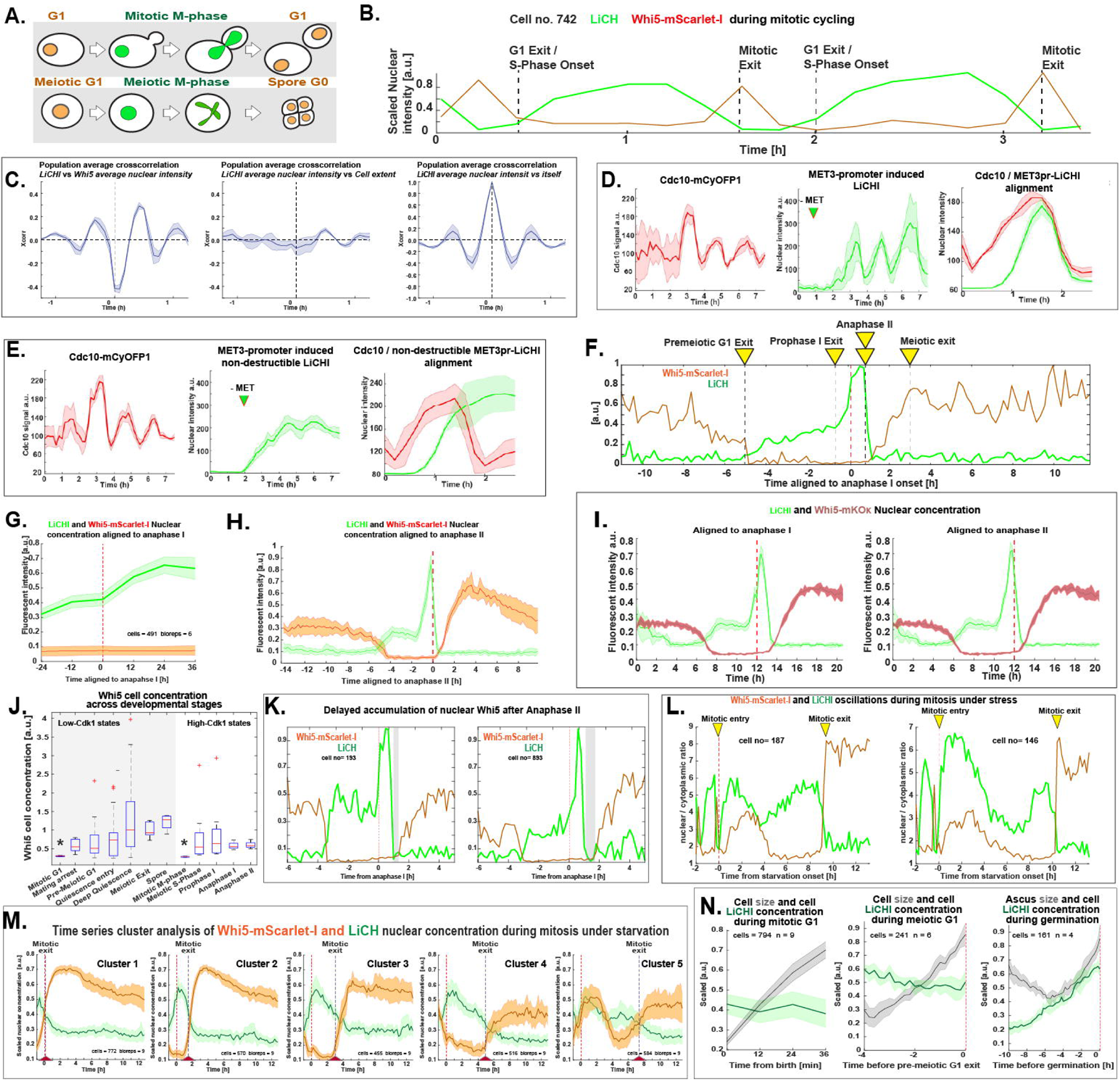
Continuous tracking of Cdk1-states across sexual life cycle stages. **(A)** Schematic comparison of the expected nuclear fluorescent signals during mitosis and meiosis using the LiCHI/Whi5 system **(B)** Representative scaled time series with labeled proliferation landmarks in a diploid cell bearing the LiCHI/Whi5 fluorescent sensor system **(C)** Crosscorrelation analysis of Whi5-mSC and LiCHI nuclear concentration time series during proliferation. Left, average crosscorrelation between Whi5-mSC and LiCHI concentration, middle, average crosscorrelation between LiCHI nuclear concentration and cell extent as negative control, right, average crosscorrelation between LiCHI nuclear concentration and itself as positive control **(D-E)** Analysis of *mNeonGreen-Ase1(aa632-885)* APC/C-Cdh1 dependent degradation in cells bearing the M-phase cell cycle marker Cdc10-mCyOFP1. Cells expressed the **(D)** *WT* or the **(E)** KEN-motif mutant of *mNeonGreen-Ase1(aa632-885*) under the MET3 promoter control. Left, aligned Cdc10 time series, middle methionine-controlled accumulation of **(D)** *WT* or the **(E)** KEN-motif mutant of *mNeonGreen-Ase1(aa632-885*); right, aligned plot of Cdc10-mCyOFP1 and *mNeonGreen-Ase1(aa632-885*) levels during the first division after *mNeonGreen-Ase1(aa632-885*) induction. Notice how the **(D)** WT but not the **(E)** KEN-motif mutant, is destroyed simultaneously with the Cdc10 signal during mitotic exit. **(F)** Representative scaled single cell LiCHI and Whi5 nuclear concentration during meiosis and sporulation with labeled landmarks. **(G)** Average Whi5-mSC and LiCHI nuclear concentration between meiotic nuclear divisions aligned to anaphase I. **(H)** Scaled average Whi5-mSC and LiCHI nuclear concentration during sporulation aligned to anaphase II **(I)** Scaled average Whi5-mmKOκ and LiCHI nuclear concentration during sporulation aligned to (left) anaphase I or (right) anaphase II. **(J)** Boxplot comparison of Whi5-mSC total cell concentration across life cycle stages identified using LiCHI/Whi5 fluorescent sensor system. **(K)** Representative scaled time series for LiCHI and Whi5 nuclear concentration aligned to anaphase I and displaying as gray area the interval with no detectable LiCHI or Whi5 signal during meiotic exit. **(L)** Representative scaled time series for LiCHI and Whi5 nuclear concentration oscillations in cells exposed to starvation (dotted line) during entry into mitotic M-phase. **(M)** Average Whi5-mSC and LiCHI nuclear concentration during the transition from proliferation into starvation (dotted line) in cells clustered according to nuclear Whi5 concentration. **(N)** Scaled average nuclear LiCHI concentration and size during exit from non-proliferative states aligned to (left) cell birth during exit from G1, (middle) pre-meiotic G1 exit, or (right) germination during the exit from the G0 spore state. Solid lines with shaded area = average ± 95% confidence intervals. Box plots display data from biological replicates: central mark, median; box bottom and top limit, 25th and 75th percentiles; whiskers, most extreme nonoutlier values. Asterisks = p > 0.05, KS test, n >3.

**Fig S6.**
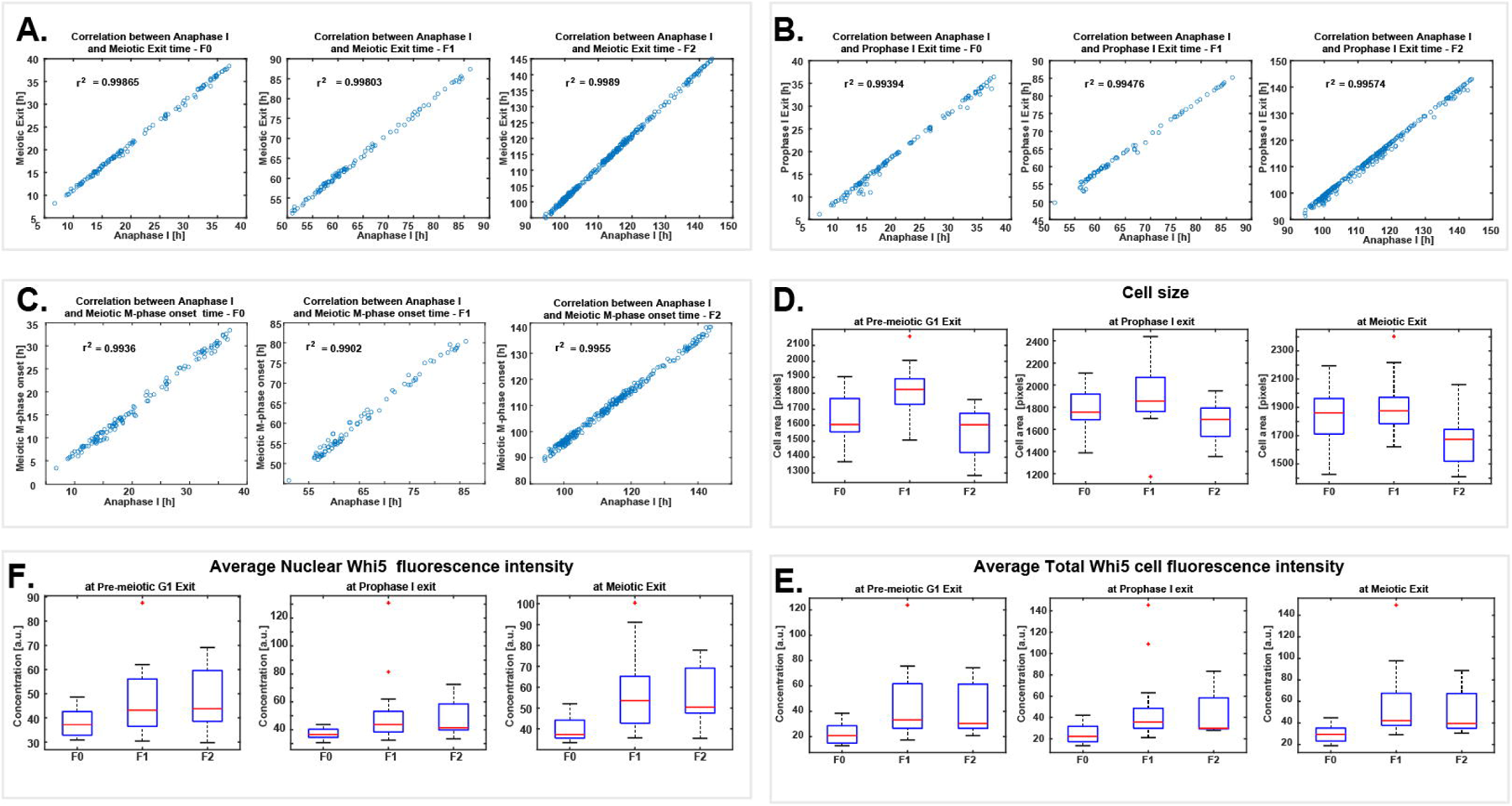
Continuous imaging of cell growth and cell division across sexually-reproducing yeast generations. **(A)** Correlation between the time of anaphase I and meiotic exit in F0, F1, and F2 generations. **(B)** Correlation between the time of anaphase I and prophase I exit in F0, F1, and F2 generations. **(C)** Correlation between the time of anaphase I and meiotic m-phase onset, defined as the point when the LiCHI signal reaches its first plateau during meiosis, in F0, F1, and F2 generations. **(D)** Boxplot comparison of cell size during pre-meiotic G1 exit, prophase I exit, and meiotic exit, in F0, F1, and F2 generations. **(E)** Boxplot comparison of average nuclear Whi5 fluorescence intensity during pre-meiotic G1 exit, prophase I exit, and meiotic exit, in F0, F1, and F2 generations. **(F)** Boxplot comparison of average total Whi5 cell fluorescence intensity during pre-meiotic G1 exit, prophase I exit, and meiotic exit, in F0, F1, and F2 generations.

